# Virus-induced vesicular acidification enhances HIV immune evasion

**DOI:** 10.1101/2024.12.17.628989

**Authors:** Marianne E. Yaple-Maresh, Giselle G. Flores, Gretchen E. Zimmerman, Cuie Chen, Kathleen L. Collins

**Author notes:** Current Address Corewell Health, St. Joseph, MI. Current Address Sysmex America, Lincolnshire, IL.

## Abstract

To block endocytic viral entry, cells typically acidify endosomes via upregulated expression of the short isoform of human nuclear receptor 7 (NCOA7) which boosts vacuolar ATPase (V-ATPase) activity. In our study, primary T cells infected with HIV-1 triggered endosomal acidification, yet NCOA7 levels were only modestly altered. Instead, we observed a pronounced depletion of the 50 kDa form of the sodium/hydrogen exchanger 6 (NHE6). Remarkably, NHE6 overexpression or treating cells with low-dose concanamycin A, a V-ATPase inhibitor, selectively neutralized endosomal pH. This neutralization impaired Nef-driven MHC-I downmodulation by our wildtype HIV reporter virus. Mechanistically, NHE6 overexpression disrupted Nef-mediated MHC-I loss by reducing recruitment of Nef to recycling endosome (Rab11^+^) compartments and blocking Nef interactions with β-COP, and ARF-1. Together, these findings reveal NHE6 as a critical regulator of endosomal pH and HIV immune evasion.

## INTRODUCTION

The pH of endosomal compartments plays a crucial role in host-pathogen interactions. For pathogens such as influenza, which enter cells via endosomes, acidic pH triggers conformational changes that promote viral fusion and entry of the viral capsid into the cytosol^1^. However, enhanced endosomal acidification can also serve as a protective mechanism against viral infection. for instance, interferon is upregulated in response to viral pathogens and induces human nuclear receptor coactivator 7 (NCOA7), which interacts with the proton pump vacuolar ATPase (V-ATPase), promoting vesicle acidification to such an extent that it inhibits influenza and severe acute respiratory syndrome (SARS) coronavirus infection^2,3^. In contrast to influenza and coronavirus, HIV enters through the cell membrane rather than endosomes, and thus its entry is not substantially modulated by changes in pH^4^. It remains unclear whether expression of NCOA7 or other proteins that influence V-ATPase activity is altered following HIV infection.

In addition to NCOA7 and V-ATPase, endosomal pH is also regulated by members of the sodium/hydrogen exchanger (NHE) family, which fine-tune pH by exchanging protons for sodium ions^5^. NHE family members selectively localize to specific cellular compartments, enabling compartment-specific regulation of pH^5^. Overexpression of NHEs can be employed as a tool to increase pH within targeted compartments and to identify the locations of pH-dependent steps in cellular trafficking pathways^6,7^.

While entry of wildtype HIV is unlikely to be inhibited by changes in pH, research utilizing concanamycin A (CMA), a V-ATPase inhibitor that disrupts acidification, suggests there may be pH sensitive steps in HIV-1 immune evasion pathways^8^. HIV-1 evades recognition by anti-HIV cytotoxic T lymphocytes (CTLs) through disruption of antigen presentation by MHC-I. This mechanism depends on the viral protein Nef, which binds to MHC-I and recruits host intracellular trafficking proteins (AP-1, ARF-1 and β-COP). These proteins target Nef-MHC-I complexes into the endolysosomal pathway from the *trans-*Golgi network (TGN), accelerating their transit to lysosomes^9–11^. Recently, we reported that CMA potently inhibits Nef-dependent MHC-I trafficking and sensitizes HIV-infected primary T cells to anti-HIV CTLs ^8^.

Although CMA disrupts lysosome acidification by inhibiting V-ATPase, our previous research demonstrated that the very low concentrations required to inhibit Nef do not affect lysosomal pH. This led us to investigate whether the pH of prelysosomal compartments is altered, and if these changes influence Nef’s ability to traffic MHC-I to the lysosome. Our data reveal that HIV infection alters the expression of proteins regulating endosomal acidity in primary T cells, specifically NCOA7 expression was modestly increased, while the 50 kDa form of NHE6, which localizes to recycling endosomes (REs), was dramatically decreased. These expression changes correlated with enhanced acidification of transferrin^+^ endosomal compartments.

Strikingly, overexpression of NHE6 in HIV-infected primary T cells partially neutralized the acidification of transferrin^+^ endosomal compartments and disrupted Nef-mediated downmodulation of MHC-I. Similarly, low concentrations of V-ATPase inhibitors that reversed Nef-dependent MHC-I downmodulation also neutralized these compartments. Mechanistic studies revealed that these pH changes disrupted localization of Nef and complex formation with its binding partners in Rab11^+^ RE compartments.

Taken together, our findings establish endosomal acidification as a pivotal regulator of Nef-driven immune evasion and underscore endosomal pH modulation as a promising strategy for counteracting HIV’s suppression of MHC-I.

## RESULTS

### HIV-1 infection leads to striking reductions in the 50 kDa form of NHE6 expression, and enhanced acidification of endosomal compartments

As part of an innate immune response to certain viral infections, levels of the V-ATPase interacting protein, NCOA7, are increased, which enhances V-ATPase activity and lowers the pH of endocytic vesicles^12^. The resulting increase in acidity is inhibitory to influenza and SARS coronavirus entry through this route^2^. Whether other viruses, such as HIV, trigger similar changes is currently unknown. In addition to V-ATPase, endosomal pH is regulated by sodium/hydrogen exchangers (NHEs) that release protons in the opposite direction of V-ATPase. To assess the impact of HIV infection on these proteins, we performed western blot analysis of primary T cells transduced with a VSV-G pseudotyped replication defective HIV construct expressing a GFP marker (Figure 1A, top panel). Because the HIV *nef* gene is a key pathogenic factor that affects intracellular trafficking in virally infected cells, we examined infection both with and without Nef expression (Nef-negative mutant referred to as ΔGPEN). For these experiments, we separated the infected (GFP^+^) from the uninfected (GFP^-^) cell subsets by fluorescence-activated cell sorting (FACS) (Figure 1B and Supplemental Figure S1A) prior to western blot analysis. As shown in Figure 1C and summarized in Figure 1E we observed a small but statistically significant increase in NCOA7 protein expression in the GFP^+^ subset of the wildtype virus that was not altered by Nef expression.

**Figure 1.**
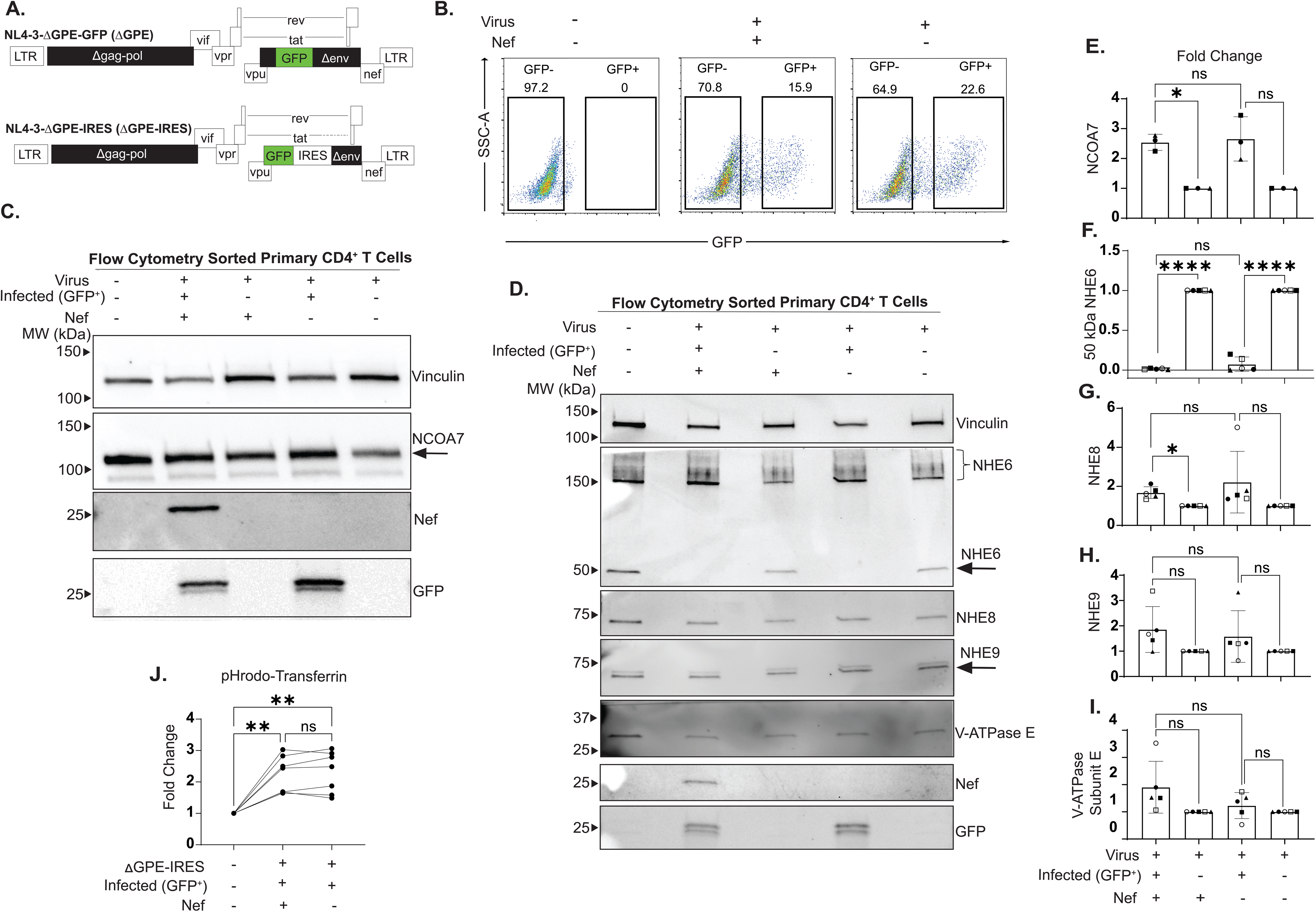
HIV-1 infection leads to striking reductions in the 50 kDa form of NHE6 expression, and enhanced acidification of intracellular compartments. (A) Schematic representation of viral genomes. Deleted genes are in black, HIV genes are in white, reporter genes are in green. (B) Representative flow plots of sorted primary CD4^+^ T cells transduced with ΔGPE or ΔGPEN. GFP+ (transduced) and GFP– (untransduced) cells were sorted by fluorescence-activated cell sorting (FACS). Mock transduced samples were used as a control to set gates. See also Supplemental Figure S1A. (C) Lysates of sorted cells from (B) were subjected to western blot analysis. (D) Western blot analysis of lysates of primary CD4^+^ T cells transduced with ΔGPE or ΔGPEN and sorted to isolate transduced and untransduced cells. See also Supplemental Figure S1B and C. (E) Summary graph of NCOA7 protein expression fold change of GFP+ and GFP–cells. Protein band intensities were quantified for 3-5 donors and normalized to vinculin. Fold change was then calculated by dividing the GFP+ vinculin-normalized signals by that of GFP-. (F) Summary graph of the 50 kDa form of NHE6 protein expression fold change of GFP+ and GFP-cells. Fold change was calculated as described in (E). (G) Summary graph of NHE8 protein expression fold change of GFP+ and GFP-cells. Fold change was calculated as described in (E). (H) Summary graph of NHE9 protein expression fold change of GFP+ and GFP-cells. Fold change was calculated as described in (E). (I) Summary graph of V-ATPase protein expression fold change of GFP+ and GFP-cells. Fold change was calculated as described in (E). (J) Summary graph of fold change of transferrin^+^ compartmental acidity in primary CD4^+^ T cells transduced with ΔGPE or ΔGPEN. Acidity of transferrin^+^ compartments in GFP+ (transduced) or mock transduced cells was evaluated by staining cells with pHrodo red-transferrin and AF-647-transferrin and subjecting to flow cytometry analysis. Fold change of pHrodo-red median fluorescence intensities (MFIs) normalized to AF-647 MFIs (to account for differences in transferrin uptake) was calculated relative to mock transduced cells and graphed. 7 donors were evaluated. Each line represents one donor. Statistical significance for (E) – (I) was determined with a One-Way ANOVA mixed effects model with Dunnett correction. Statistical significance for (J) was determined with a One-Way ANOVA mixed effects model with Tukey correction. * p < 0.05, ** p < 0.01, *** p < 0.001, **** p < 0.0001.

Because sodium/hydrogen exchangers oppose V-ATPase activity to increase the pH of endosomal compartments, we examined expression of NHE6, NHE8 and NHE9, which localize to different intracellular compartments. Prior experiments performed in various cell types revealed that NHE6 localizes to the RE^5,13^, NHE8 to the *mid* and *trans*-Golgi^5^, and NHE9 to the late endosome^5^. Consistent with published results, NHE8 and NHE9 migrate as a single species at approximately 75 kDa (Figure 1D). In contrast, NHE6 migrates as multiple forms that include high molecular weight oligomers and aggregates as well as smaller, 50 and/or 75 kDa forms that can vary with the cell type ^14–17^. Given this complexity, we confirmed that the bands labeled as NHE6 in Figure 1D were bona fide forms of the protein by using the NHE6 peptide epitope to competitively inhibit specific antibody binding (Supplemental Figure S1B and C).

Surprisingly, following exposure to HIV (with or without Nef), infected (GFP^+^) cells had approximately 65-fold less of the 50 kDa form of NHE6 (Figure 1D, summarized in 1F). In contrast, levels of NHE8 increased slightly but significantly (Figure 1D, summarized in Figure 1G) and NHE9 levels did not significantly change (Figure 1D, summarized in 1H). Levels of V-ATPase subunit E also did not significantly change in response to HIV infection (Figure 1D, summarized in Figure 1I).

The observation that NHE6 expression decreased while NCOA7 levels slightly but significantly increased upon infection led us to hypothesize that the pH of endosomal compartments may become more acidic following HIV-1 infection. To interrogate this, we utilized a pH-sensitive conjugate of transferrin (pHrodo-transferrin), which is endocytosed and delivered to the RE before being trafficked back to the cell surface^18^. To establish transferrin’s localization in T cells, we examined its distribution relative to Rab11, a marker of the RE, Rab5, an early endosomal marker, and overexpressed NHE6. Under our assay conditions, transferrin showed substantial overlap with both Rab11 and Rab5 (Supplemental Figure S1D and quantified in S1E). Additionally, we observed colocalization between transferrin and overexpressed NHE6 (Supplemental Figure S1D), supporting the use of pHrodo-transferrin to assess the effect of NHE6 overexpression on endosomal pH.

To control for potential differences in the amount of pHrodo-transferrin taken up by the cells upon infection, we also included transferrin conjugated to AlexaFluor 647 (AF-647), a dye that is not known to have pH-dependent properties. Thus, changes in transferrin^+^ compartmental pH could be monitored by normalizing the pHrodo-transferrin signal to the AF-647-transferrin signal.

Consistent with our hypothesis, we found that transferrin^+^ compartments from infected (GFP^+^) primary T cells were significantly more acidic than those from mock infected cells (Figure 1J). Consistent with our western blot findings that infection alters NCOA7 and NHE6, we observed that infection-induced acidification was unaffected by Nef expression. Together, these data suggest the hypothesis that the HIV-1-induced decrease in the 50 kDa form of NHE6 contributed to the decrease in pH of endosomal compartments. Moreover, based on colocalization of transferrin, Rab11 and NHE6 (Supplemental Figure S1D), REs are likely to be amongst the most impacted compartments.

### Overexpression of endosomal sodium/hydrogen exchangers reduces the ability of HIV-1 Nef to disrupt MHC-I trafficking

Although HIV entry is unlikely to be influenced by endosomal acidification, unlike other viruses, Nef exploits these compartments to direct MHC-I to the lysosome. The pronounced decrease in expression of the 50 kDa form of NHE6, accompanied by increased acidification of transferrin^+^ compartments in HIV-infected primary T cells led us to investigate whether the pH of these compartments affects Nef’s ability to traffic MHC-I to the lysosome. To address this, we generated HIV-1 constructs that overexpress NHE6 or NHE8 using an internal ribosome entry site (IRES, referred to as ΔGPE-IRES-NHE6/8). Successful overexpression of the NHEs following viral transduction was confirmed with fluorescence confocal microscopy (Figure 2A-D), with additional validation by western blot (Supplemental Figure S2A and S2B). We found that overexpression of both NHEs significantly reduced Nef-dependent MHC-I downmodulation (Figure 2E and F). Notably, NHE6 overexpression decreased MHC-I downmodulation to levels comparable to those observed with a Nef negative mutant (ΔGPEN). Overexpression of NHEs also inhibited Nef-dependent CD4 downmodulation, but this effect was only significant for NHE6, and CD4 downmodulation remained higher than in cells transduced with the Nef-negative mutant virus (Figure 2G). Thus, the impact of NHE overexpression appears to be greater for MHC-I than for CD4.

**Figure 2.**
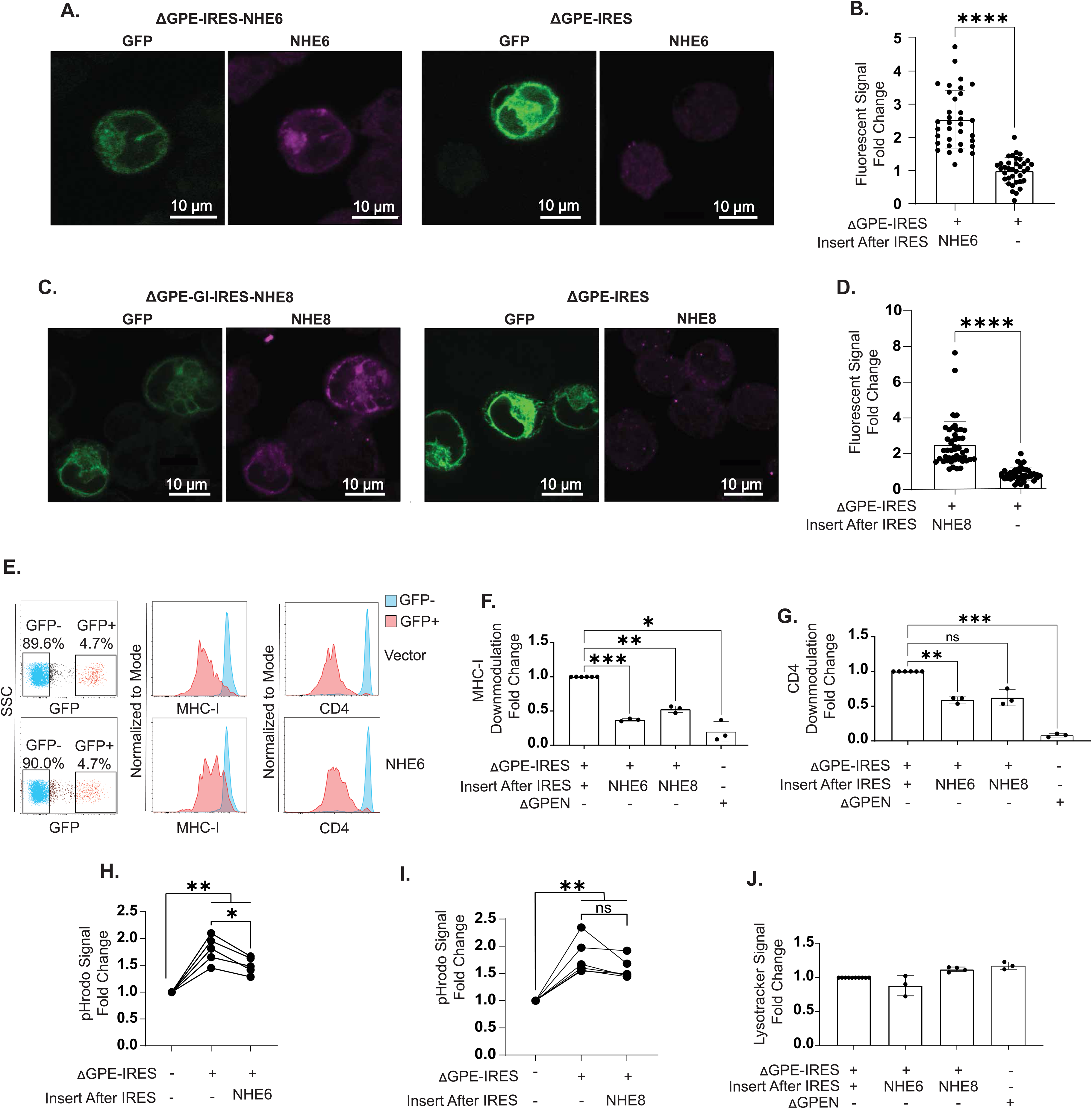
Overexpression of endosomal sodium/hydrogen exchangers reduces the ability of HIV-1 Nef to disrupt MHC-I trafficking. (A) Representative fluorescence confocal microscopy images of CEM-A2 cells 48 hr after transduction with the indicated viruses and stained for NHE6. Identical laser settings were used for all images of each experimental replicate. See also Supplemental Figure S2A (B) Summary graph of NHE6 expression 48 hr post transduction of CEM-A2 cells with the indicated viruses. Corrected total cell fluorescence (CTCF) was calculated for each cell. Fold change in NHE6 expression was calculated by dividing the CTCF of all cells by the average ΔGPE-IRES CTCF. At least 30 cells were imaged for each condition. (C) Representative images of CEM-A2 cells 48 hr after transduction with the indicated viruses and stained for NHE8. See also Supplemental Figure S2A and B. (D) Summary graph of NHE8 expression 48 hr post transduction of CEM-A2 cells with the indicated viruses. Fold change was calculated as described in (B). (E) Representative flow plots of primary CD4^+^ T cells stained for surface MHC-I or CD4 expression 72 hr post transduction with the indicated viruses. (F) Summary graph of surface MHC-I expression in primary CD4^+^ T cells 72 hr post transduction with the indicated viruses. MHC-I MFIs were obtained for GFP+ and GFP-cells. MHC-I downmodulation was calculated by dividing MHC-I MFIs of GFP+ cells by GFP-cells. The fold change was calculated by dividing the resulting values by that of the control virus (ΔGPE-IRES). 3 donors were evaluated. (G) Summary graph of Nef-dependent CD4 downmodulation in primary CD4^+^ T cells 72 hr post transduction with the indicated viruses. CD4 downmodulation and fold change was calculated as described in (F), except CD4 MFIs were assessed. 3 donors were evaluated. (H) Summary graphs of transferrin^+^ compartmental acidity of primary CD4^+^ T cells 72 hr post transduction with the indicated viruses. Cells were stained with pHrodo-transferrin and AF-647-transferrin. Fold change in pHrodo signal was evaluated as described in Figure 1J. 5 donors were evaluated. Each line represents 1 donor. (I) Summary graphs of transferrin^+^ compartmental acidity of primary CD4^+^ T cells 72 hr post transduction with the indicated viruses. Fold change was calculated as described in Figure 1J. 6 donors were evaluated. Each line represents 1 donor. (J) Summary graph of lysosome acidification in primary CD4^+^ T cells 72 hr post transduction with the indicated viruses. Fold change was determined by dividing Lysotracker Red MFIs from GFP+ cells with Lysotracker Red MFIs from GFP+ cells transduced with the control virus (ΔGPE-IRES). 3 donors were evaluated. See also Supplemental Figure S2C and D. Statistical significance for (B) and (D) was determined with an unpaired T test. Statistical significance for (F) and (G) was determined with a One-way ANOVA mixed effects analysis with Dunnett correction. Statistical significance for (H) and (I) was determined with a One-Way ANOVA mixed effect analysis with Šidák correction. * p < 0.05, ** p < 0.01, *** p < 0.001, **** p < 0.0001.

To assess whether NHE6 overexpression was sufficient to neutralize endosomal compartments, we utilized pHrodo-transferrin. Compared to control cells transduced with an “empty” HIV IRES-containing construct, NHE6 overexpression significantly reduced acidification of transferrin^+^ compartments (Figure 2H), partially reversing HIV-induced endosomal acidification. In contrast, overexpression of NHE8 did not significantly counteract HIV-induced endosomal acidification (Figure 2I), consistent with its primary localization to the Golgi. Notably, NHE6 overexpression in the absence of HIV infection did not significantly neutralize transferrin^+^ compartments or alter surface MHC-I expression (Supplemental Figures S2C and D). Importantly, neither NHE6 nor NHE8 overexpression affected lysosomal pH in primary T cells (Figure 2J). Collectively, these data support our hypothesis that NHE6 selectively counteracts virus-induced endosomal acidification and disrupts Nef-dependent MHC-I downmodulation.

### Overexpressed NHE6 primarily localizes to REs

Overexpression of both NHE6 and NHE8 significantly impaired Nef-mediated downmodulation of MHC-I. While the TGN is a well-established site for Nef-directed MHC-I trafficking^9^, the role of the RE, the target compartment of NHE6, was surprising and required further validation. To accomplish this, we sought to confirm the localization of overexpressed NHE6 and NHE8 in CEM-A2 T cells. Using fluorescence confocal microscopy, we verified that NHE8 overexpression in cells transduced with ΔGPE-IRES-NHE8 indeed localized predominantly to the TGN, as indicated by colocalization with the TGN marker AP-1 γ (Figure 3A and Supplemental Figure S3A, quantification in Supplemental Figure S3B). Because AP-1 is known to interact with Nef, we also confirmed that AP-1 staining patterns in mock infected cells were comparable to those in cells transduced with the Nef-expressing viral construct (Supplemental Figure S3A).

**Figure 3.**
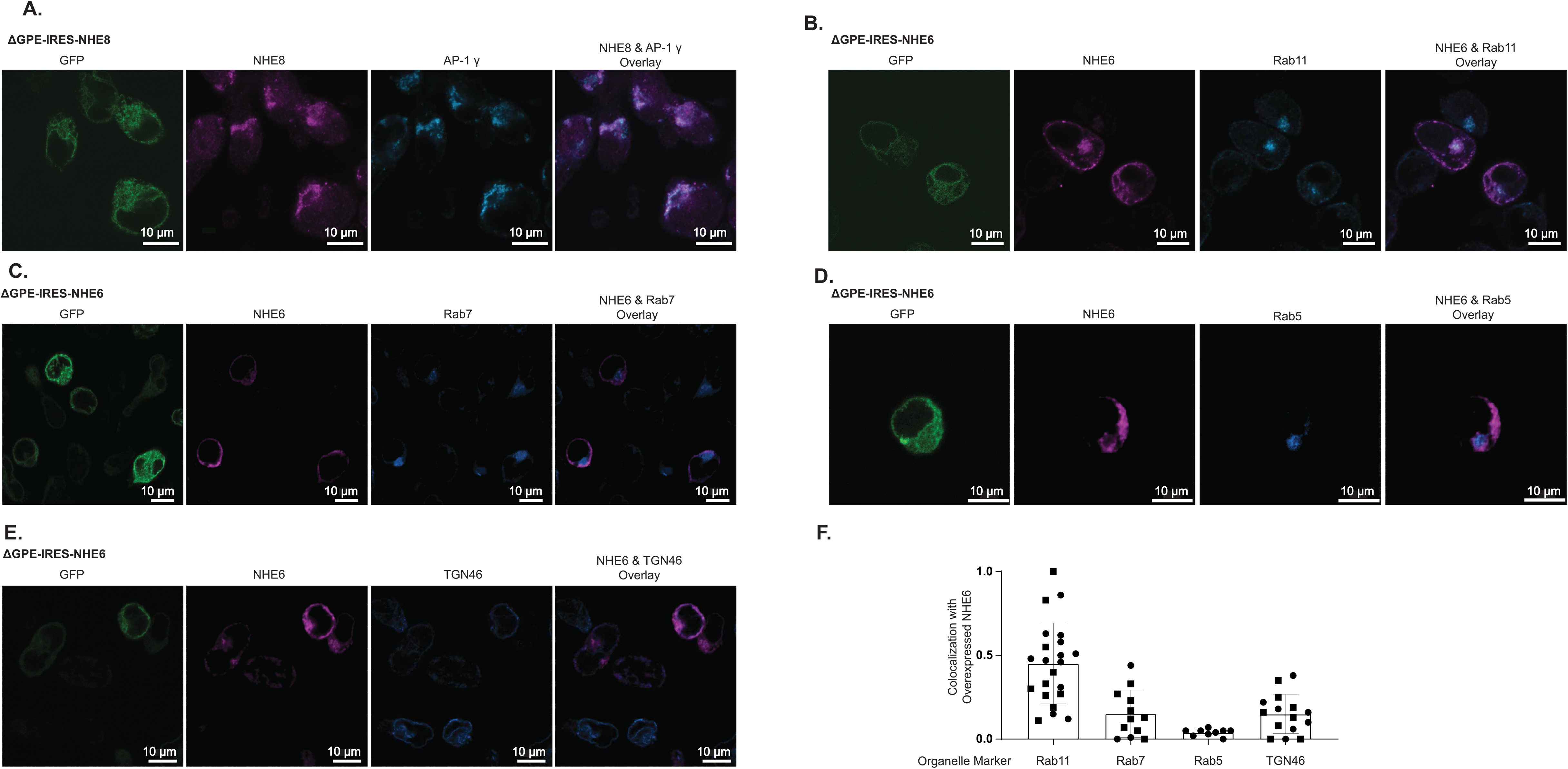
Overexpressed NHE6 Primarily localizes to REs. (A) Representative fluorescence confocal microscopy images of CEM-A2 cells 48 hr after transduction with the indicated virus and stained for NHE8 and AP-1 γ. Identical laser settings were used for all images of each experimental replicate. See also Supplemental Figure S3A and B. (B) Representative fluorescence confocal microscopy images of CEM-A2 cells 48 hr after transduction with the indicated virus and stained for NHE6 and Rab11. See also Supplemental Figure S3C. (C) Representative fluorescence confocal microscopy images of CEM-A2 cells 48 hr after transduction with the indicated virus and stained for NHE6 and Rab7. See also Supplemental Figure S3D. (D) Representative fluorescence confocal microscopy images of CEM-A2 cells 48 hr after transduction with the indicated virus and stained for NHE6 and Rab5. See also Supplemental Figure S3E. (E) Representative fluorescence confocal microscopy images of CEM-A2 cells 48 hr after transduction with the indicated virus and stained for NHE6 and TGN46. See also Supplemental Figure S3F. (F) Summary graph of quantification of colocalization of overexpressed NHE6 with the indicated organelle markers. Colocalization was quantified using Imaris software and creating spot masks for NHE6 and the organelle staining. The average diameter of puncta for each stain was used to create spots. Spots were considered colocalized if the distance between them was less than or equal to the sum of the radii of the individual spots. The number of NHE6 spots colocalized with the indicated organelle marker was divided by the total number of NHE6 spots and graphed. Each point represents quantification of 1 image. Symbols represent biological replicates. 1-2 biological replicates per staining were conducted. The number of cells quantified per image ranged from 1-8 cells. The total number of images analyzed per staining ranged from 12-31. All organelle markers were imaged in the TRITC channel and pseudocolored blue for easier visualization. See also Supplemental Figure S4.

We further demonstrated that overexpressed NHE6 predominantly colocalizes with Rab11, a marker of the RE (Figure 3B; quantified in 3F). In addition, we confirmed that NHE6 showed minimal colocalization with markers of the late endosome (Rab7), early endosome (Rab5), or the TGN (TGN46) (Figure 3C-E; quantified in Figure 3F and detailed in Supplemental Figures S3C-F). Additionally, overexpressed NHE6 localized to Rab11^+^ compartments in a pattern similar to endogenous NHE6 (Supplemental Figure S3C).

The preferential localization of overexpressed NHE6 to Rab11^+^ compartments, rather than Rab5^+^ compartments contrasts with that of transferrin, which distributes relatively evenly between both (Supplemental Figures S1D and S1E). The precise targeting of NHE6 to Rab11^+^ compartments allows us to conclude that the change in pHrodo-transferrin signal observed upon NHE6 overexpression (Figure 2H) was primarily due to the neutralization of Rab11^+^ compartments by NHE6.

We also sought to assess Nef’s requirement for late endosomal acidification by overexpressing NHE9. Although we generated a viral construct that successfully overexpressed NHE9, we were unable to confirm its localization to the late endosome in T cells (Supplemental Figure S4). Consequently, it was excluded from further analysis.

### V-ATPase inhibitors neutralize transferrin^+^ compartmental pH at concentrations that reverse Nef activity

We previously reported that V-ATPase inhibitors, such as CMA, inhibit Nef-dependent MHC-I downmodulation^8^. Building on the observation that reducing endosomal acidity via NHE6 overexpression blocks Nef-mediated MHC-I downmodulation, we investigated whether CMA neutralizes endosomal compartments at concentrations effective against Nef. As shown in Figure 4A, CMA reversed Nef-dependent MHC-I downmodulation (“Nef activity”) at remarkably low concentrations (IC_50_ ∼0.1 nM), which are well below those required to impact lysosomal acidification (IC_50_ ∼1 nM) in primary T cells transduced with an NL4.3 ΔGPE reporter virus harboring FLAG-tagged Nef. Notably, the IC_50_ for CMA-induced neutralization of transferrin^+^ compartments closely matched that for reversal of Nef activity (Figure 4B), and both were significantly lower than the concentration needed for lysosome neutralization (Figure 4C).

**Figure 4.**
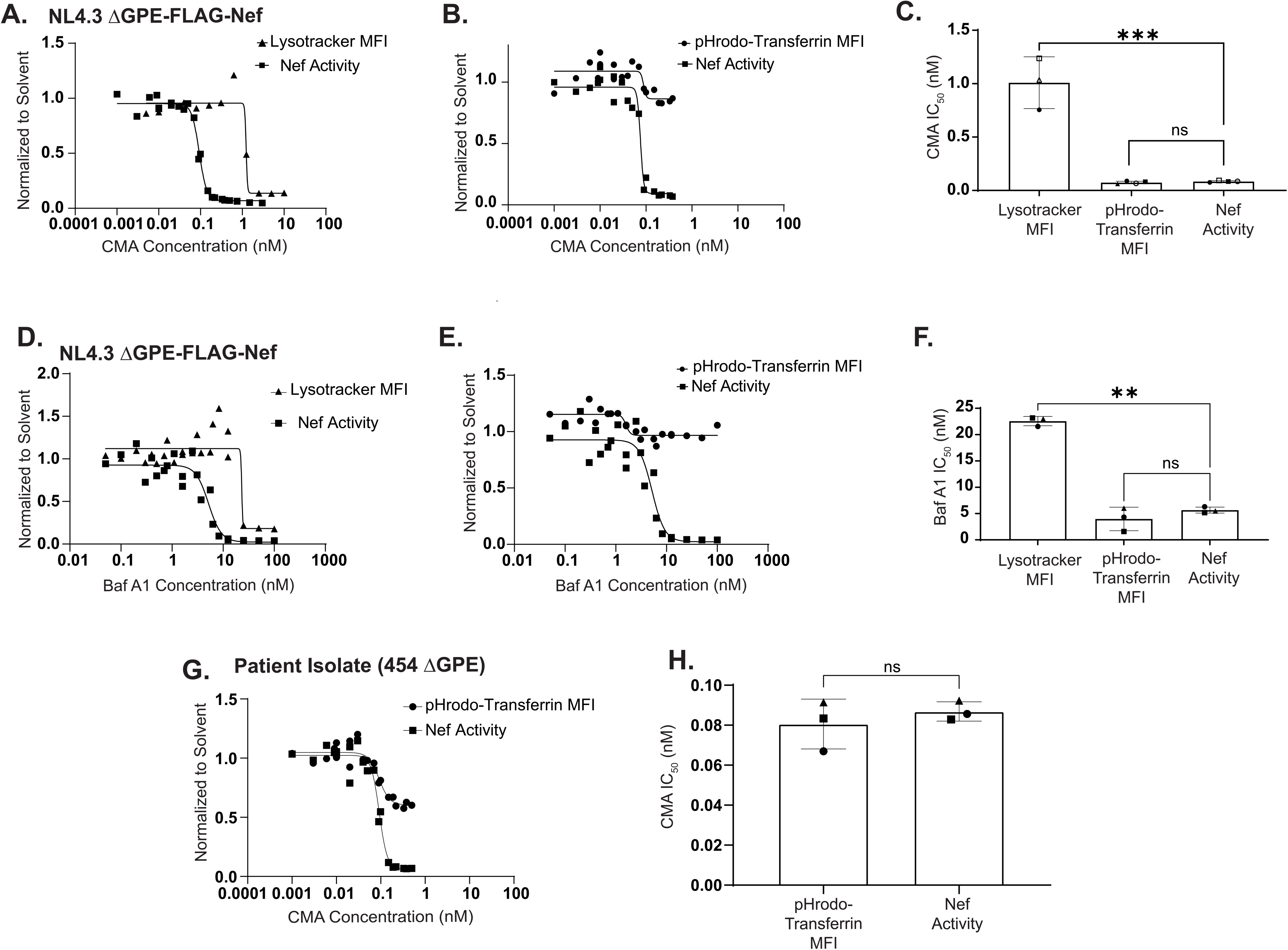
V-ATPase inhibitors neutralize transferrin+ compartmental pH at concentrations that reverse Nef activity. (A) CMA dose response curves for primary CD4^+^ T cells transduced with ΔGPE virus harboring FLAG-tagged Nef (ΔGPE-FLAG-Nef) and treated with CMA or solvent two days after transduction for 24 hr. CMA concentrations ranged from 10 – 0.005 nM (Lysotracker MFI) or 3 – 0.001 nM (Nef Activity). (B) CMA dose response curves for primary CD4^+^ T cells transduced with ΔGPE-FLAG-Nef and treated with CMA or solvent 48 hr after transduction for 24 hr then stained with pHrodo and AF-647 transferrin. CMA concentrations ranged from 0.38 – 0.001 nM. (C) Summary graphs of CMA IC_50_s for Lysotracker MFI, pHrodo-transferrin MFI, and Nef activity. (D) Bafilomycin A1 (Baf A1) dose response curves for primary CD4^+^ T cells transduced with ΔGPE-FLAG-Nef and treated with Baf A1 or solvent two days after transduction for 24 hr. Baf A1 concentrations ranged from 100 – 0.05 nM. (E) Baf A1 dose response curves for primary CD4^+^ T cells transduced with ΔGPE-FLAG-Nef and treated with Baf A1 or solvent two days after transduction for 24 hr then stained with pHrodo and AF-647 transferrin. Baf A1 concentrations ranged from 100 – 0.05 nM. (F) Summary graphs of Baf A1 IC_50_s for Lysotracker MFI, pHrodo-transferrin MFI, and Nef activity. (G) CMA dose response curves for primary CD4^+^ T cells transduced with 454 ΔGPE and treated with CMA or solvent two days after transduction for 24 hr. CMA concentrations ranged from 0.5 nM – 0.001 nM. (H) Summary graphs of CMA IC_50_s for pHrodo-transferrin MFI, and Nef activity. Statistical significance for (C) and (F) was determined with a One-way ANOVA mixed effects analysis with Dunnett correction. Statistical significance for (H) was determined with a paired T test. 3 donors were evaluated. Each symbol represents one donor. * p < 0.05, ** p < 0.01, *** p < 0.001, **** p < 0.0001.

Similar findings were observed with another V-ATPase inhibitor, Bafilomycin A1 (Baf A1), which inhibits Nef at higher concentrations than CMA, but still demonstrates a substantial difference between the IC_50_ for Nef inhibition and lysosomal neutralization (Figure 4D)^8^. As with CMA, the amount of Baf A1 required to neutralize transferrin^+^ compartments was nearly identical to that needed to inhibit Nef activity (Figure 4E and 4F). Importantly, CMA also neutralized transferrin^+^ compartmental pH at concentrations that inhibited Nef activity in cells infected with a patient-derived reporter virus (Figure 4G and 4H).

Collectively, these results demonstrate that neutralization of endosomes, either through NHE6 overexpression or low-dose V-ATPase inhibitor treatment, impairs Nef’s ability to downmodulate MHC-I from the surface of infected cells.

### Neutralization of transferrin^+^ compartments via NHE6 overexpression prevents Nef from binding to cellular proteins required for Nef-mediated disruption of MHC-I trafficking

We next investigated the mechanism by which neutralization of transferrin^+^ compartments through NHE6 overexpression reversed Nef-dependent MHC-I downmodulation. Prior studies have shown that Nef is required in the TGN, where it forms a three-way complex with the MHC-I cytoplasmic tail, and clathrin adaptor protein AP-1^9,11^, a process dependent on the GTPase ARF-1^9,19^. These interactions are crucial for Nef to disrupt MHC-I trafficking from Golgi to the cell surface. Instead, Nef-bound MHC-I is redirected to less-well characterized compartments, where interactions with β-COP of the coatomer complex commit MHC-I to lysosomal degradation^11^. We hypothesized that endosomal pH neutralization via NHE6 overexpression could interfere with Nef’s ability to form these complexes with key host proteins, thereby impairing MHC-I trafficking.

To test this hypothesis, we engineered an HIV construct overexpressing NHE6 that also included Nef tagged with a FLAG epitope. We verified that insertion of the tag into the N-terminal loop did not impair Nef’s ability to downmodulate MHC-I (Figure 5A and B). Notably, FLAG-tagged Nef exhibited robust, ∼50-fold downmodulation of CD4 that was slightly less than wildtype Nef, but still highly effective (Figure 5A and B). Western blot analysis of cell lysates transduced with either the empty vector or the NHE6 overexpressing virus confirmed expression of the relevant proteins prior to FLAG bead incubation (Input; Figure 5C).

**Figure 5.**
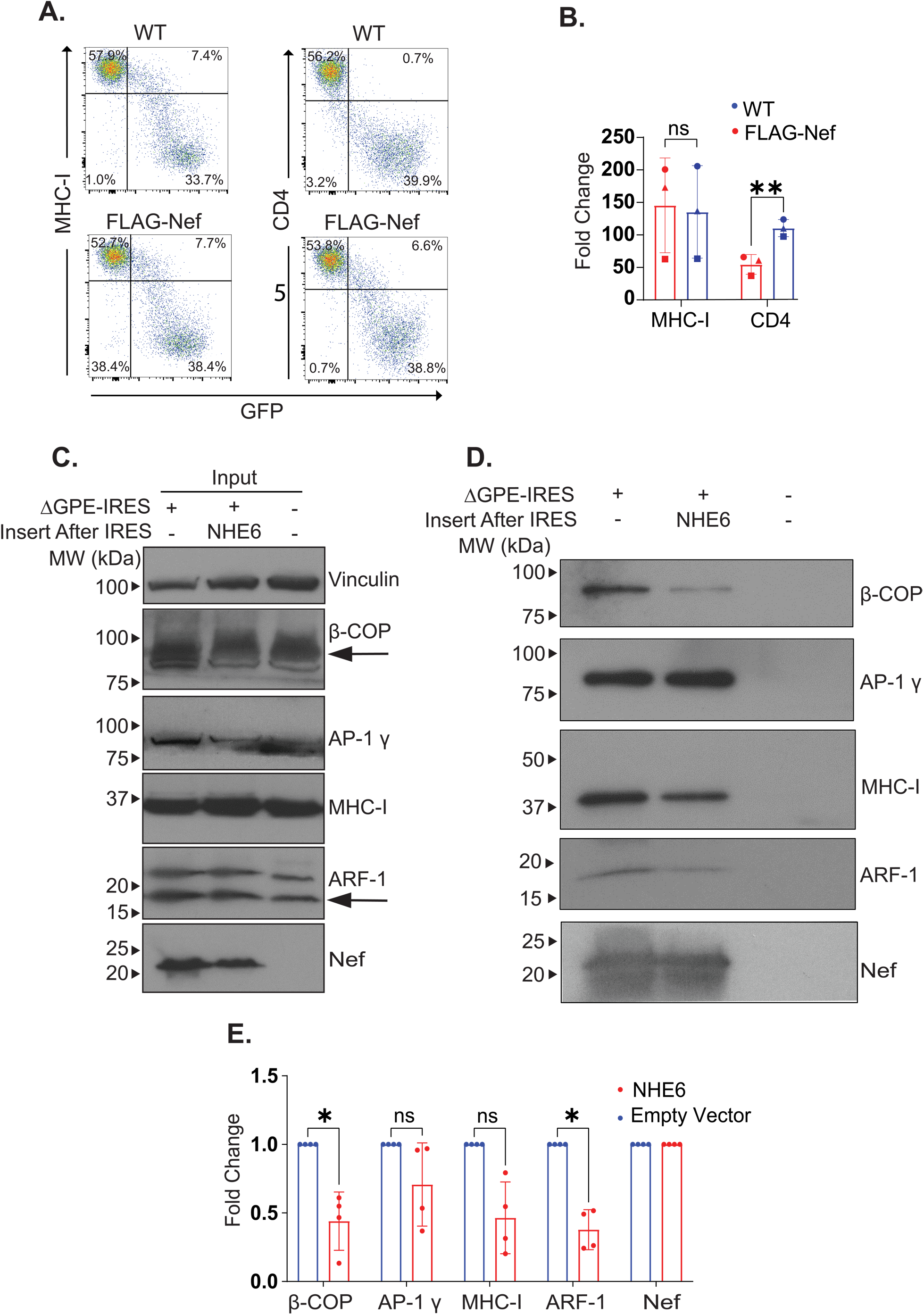
Neutralization of transferrin+ compartments via NHE6 overexpression prevents Nef from binding to cellular proteins required for Nef-mediated disruption of MHC-I trafficking. (A) Representative flow plots showing MHC-I and CD4 downmodulation in CEM-A2 cells 48 hr post transduction with ΔGPE-IRES (WT) or ΔGPE-FLAG-Nef-IRES (FLAG-Nef). (B) Summary of fold change of CD4 and MHC-I downmodulation with or without the FLAG sequence inserted in Nef. Downmodulation was calculated as described in Figure 2F. 3 experimental replicates were performed. Each symbol represents 1 replicate. (C) Representative western blot analysis of input (before incubation with FLAG beads) fractions of CEM-A2 lysates transduced with the indicated viruses. (D) Representative Co-IP western blot analysis of CEM-A2 cell lysates transduced with the indicated viruses. (E) Summary graphs of protein pulled down with Nef with or without NHE6 overexpression. Protein band signals were quantified and normalized to the Nef signal to account for slight differences in the amount of Nef pulled down between samples. Fold change of protein bound to Nef was calculated by dividing the Nef-normalized signals of NHE6-overexpressing samples by that of empty vector samples. 4 experimental replicates were performed. See also Supplemental Figure S5. Statistical significance was determined with a Paired T Test. * p < 0.05, ** p < 0.01, *** p < 0.001, **** p < 0.0001.

Furthermore, FLAG-tagged Nef expressed from the empty vector interacted as expected with β-COP, AP-1, MHC-I, and ARF-1 (Figure 5D). Remarkably, NHE6 overexpression led to selective loss of Nef interactions with β-COP and ARF-1, while Nef-AP-1 and Nef-MHC-I associations were relatively preserved (Figure 5E). Importantly, overexpression of NHE6 did not significantly alter β-COP or ARF-1 protein levels (Supplemental Figure S5).

In summary, these findings demonstrate that interactions between β-COP, ARF-1 and Nef are highly sensitive to endosomal pH changes. Moreover, they underscore the involvement of NHE6-occupied compartments, such as REs, in Nef-mediated trafficking of MHC-I to the lysosome. Because both MHC-I and CD4 downmodulation depend on Nef’s interactions with β-COP, these results explain the inhibition of MHC-I and CD4 downmodulation observed following NHE6 overexpression (Figure 2F and G).

### NHE6 overexpression disrupts recruitment of Nef to the recycling endosome

To further elucidate how reduced endosomal acidity affects Nef-dependent MHC-I downmodulation, we examined whether Nef localizes to the Rab11^+^ compartments and if this localization is pH-sensitive. Using a CD4^+^ T cell line stably expressing FLAG-tagged Rab11 (CEM-A2-FLAG-Rab11) and a previously published protocol for isolating intact Rab-positive compartments (Figure 6A)^20^, we first confirmed successful infection with reporter viruses overexpressing NHE6 (Figure 6B) and verified expression of our proteins of interest in whole cell lysates (Figure 6C). We then isolated Rab11^+^ vesicles and analyzed their protein composition via western blot analysis (Figure 6D).

**Figure 6.**
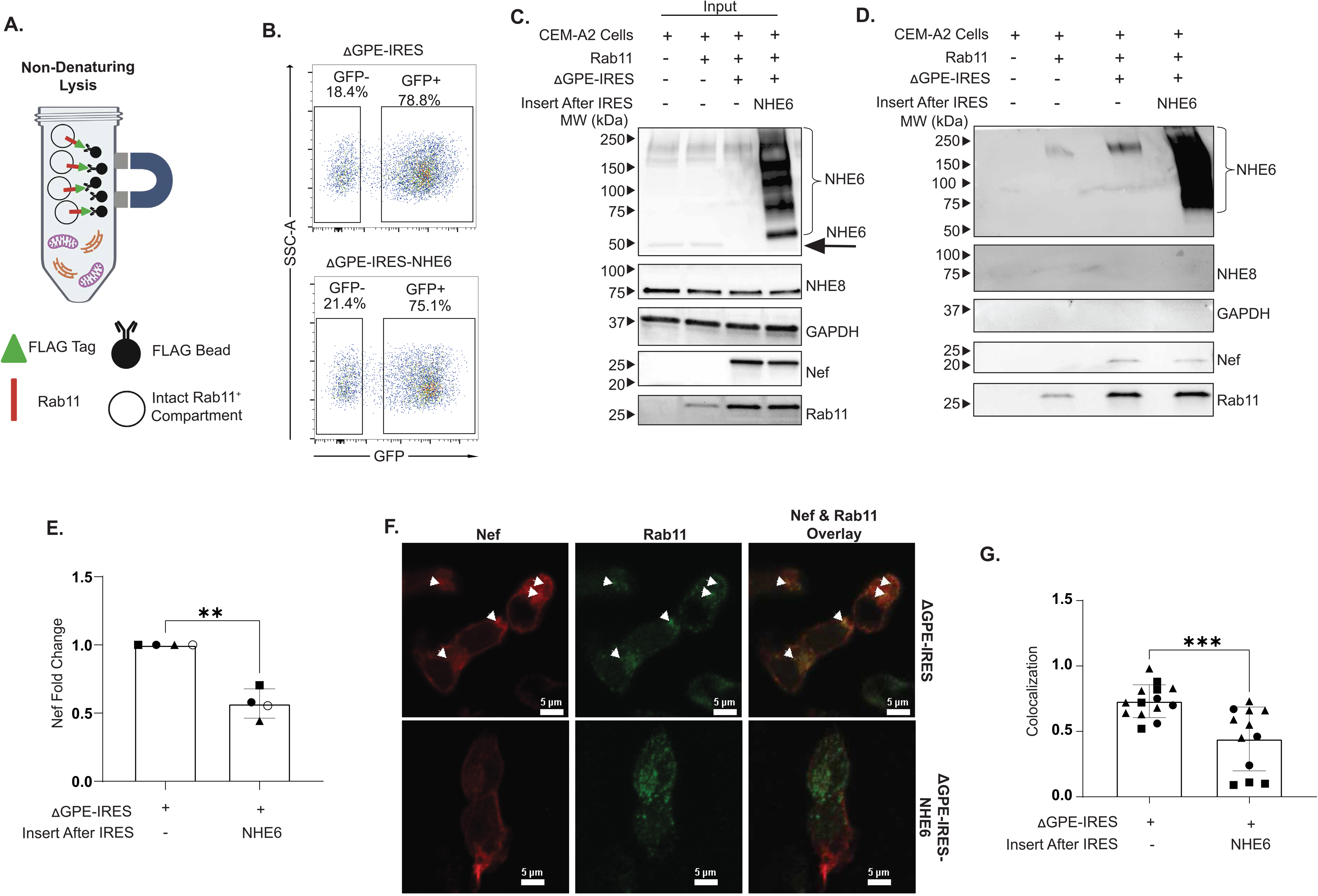
NHE6 overexpression disrupts recruitment of Nef to the recycling endosome. (A) Schematic diagram of the method to precipitate intact Rab11^+^ compartments from CEM-A2 cells stably expressing FLAG-tagged Rab11 (CEM-A2-FLAG-Rab11). (B) Representative flow plots showing transduction rates of CEM-A2-FLAG-Rab11 cells 48 hr post transduction with the indicated viruses. (C) Representative western blot analysis of input (prior to incubation with FLAG beads) fractions of lysates from CEM-A2-FLAG-Rab11 cells 48 hr post transduction with the indicated viruses. (D) Representative Co-IP western blot analysis of isolated Rab11^+^ compartments from CEM-A2-FLAG-Rab11 lysates transduced with the indicated viruses and precipitated with FLAG beads. (E) Summary graph of Nef expression in isolated Rab11^+^ compartments during HIV-1 transduction with or without NHE6 overexpression. Nef signal was normalized to the Rab11 signal to account for slight variations in the amount of Rab11 pulled down. Fold change was then calculated by dividing the resulting value for NHE6 overexpressing samples by that of the control virus (ΔGPE-IRES) samples. 4 experimental replicates were performed. Symbols represent 1 replicate. (F) Representative fluorescence confocal microscopy images of CEM-A2 cells 48 hr post transduction with the indicated viruses and stained for Nef (FLAG) and Rab11. Rab11 was imaged in the Cy5 channel and pseudocolored to green for easier visualization. See also Supplemental Figure S6. (G) Summary graph of quantification of colocalization between Nef and Rab11. Colocalization was quantified as described in 3F except a surface mask was assigned to the Nef staining to account for its diffused staining throughout the cell. Nef and Rab11 were considered colocalized if the distance between the Rab11 spot and Nef surface was less than or equal to 0. The number of Rab11 spots colocalized with Nef was divided by the total number of Rab11 spots and graphed. Each point represents 1 image. The number of cells quantified per image ranged from 1-9. Each symbol represents a biological replicate. 3 biological replicates were performed. The number of images analyzed per experiment ranged from 6-13. Statistical significance for (E) was determined with a paired T test. Statistical significance for (G) was determined with an unpaired T test. * p < 0.05, ** p < 0.01, *** p < 0.001, **** p < 0.0001.

As expected, we detected both endogenous and overexpressed NHE6 within Rab11^+^ compartments, confirming its natural localization (Figure 6D). In contrast, NHE8 and GAPDH were absent in these purified vesicles. Importantly, we observed that Nef was recruited to Rab11^+^ compartments in infected T cells (Figure 6D). Strikingly, overexpression of NHE6 markedly diminished Nef recruitment to these compartments (Figure 6D; summarized in 6E). To validate this result, we also employed confocal fluorescence microscopy to monitor Nef colocalization with Rab11^+^ compartments in T cells infected with control or NHE6-overexpressing reporter viruses (Figure 6F and Supplemental Figure S6). Consistent with our findings from Rab11^+^ vesicle isolation, confocal microscopy analysis revealed that Nef recruitment to Rab11^+^ compartments was markedly diminished following endosomal pH neutralization by NHE6 overexpression (Figure 6G).

Together, these results explain the reduced complex formation observed upon endosomal neutralization and highlight the critical role of Rab11^+^ compartments in Nef-dependent trafficking of MHC-I within HIV-infected cells.

## DISCUSSION

### HIV-induced endosomal acidification

Here we describe a link between HIV-1 induced acidification of endosomal compartments and Nef-mediated trafficking of MHC-I to the lysosome. Our findings show a dramatic, virus-driven downmodulation of the 50 kDa form of NHE6, a sodium/proton exchanger that localizes to Rab11/transferrin^+^ compartments. This loss is complemented by increased expression of NCOA7, a protein that promotes vesicular acidification by interacting with V-ATPase. Consequently, we observed a significant rise in acidity within transferrin^+^ compartments during HIV-1 infection; this acidification could be counteracted by CMA treatment or NHE6 overexpression. Acidification of endosomes is known to be protective in certain viral infections, as it blocks viral entry^12^ and enhances antigen presentation via MHC class II^21,22^. However, HIV employs a distinct strategy: while its entry is not influenced by increased endosomal acidification, the virus employs Nef to redirect MHC-I to acidified compartments, ultimately targeting it for lysosomal degradation^23–25^.

### pH-Dependency of Nef activity

Consistent with a pivotal role for endosomal acidification in Nef-mediated MHC-I trafficking, we recently reported the V-ATPase inhibitor CMA blocks Nef-dependent MHC-I downmodulation at concentrations far lower than required to impact lysosomal pH^8^. Our observation of a marked decrease in the expression of the 50 kDa form of NHE6, accompanied by increased acidification of transferrin^+^ compartments, led us to hypothesize that CMA inhibits Nef by disrupting the pH of Rab11^+^ endosomal compartments where NHE6 resides.

Supporting this hypothesis, we found that both CMA and Baf A1 neutralized transferrin-containing compartments (many of which are Rab 11^+^) at concentrations comparable to those that reversed Nef-dependent MHC-I downmodulation, and notably lower than those needed for lysosomal neutralization. These findings demonstrate that Nef activity is disrupted when endosomal acidity is reduced, and they provide new insight into the mechanism of action of CMA (Figure 7).

**Figure 7.**
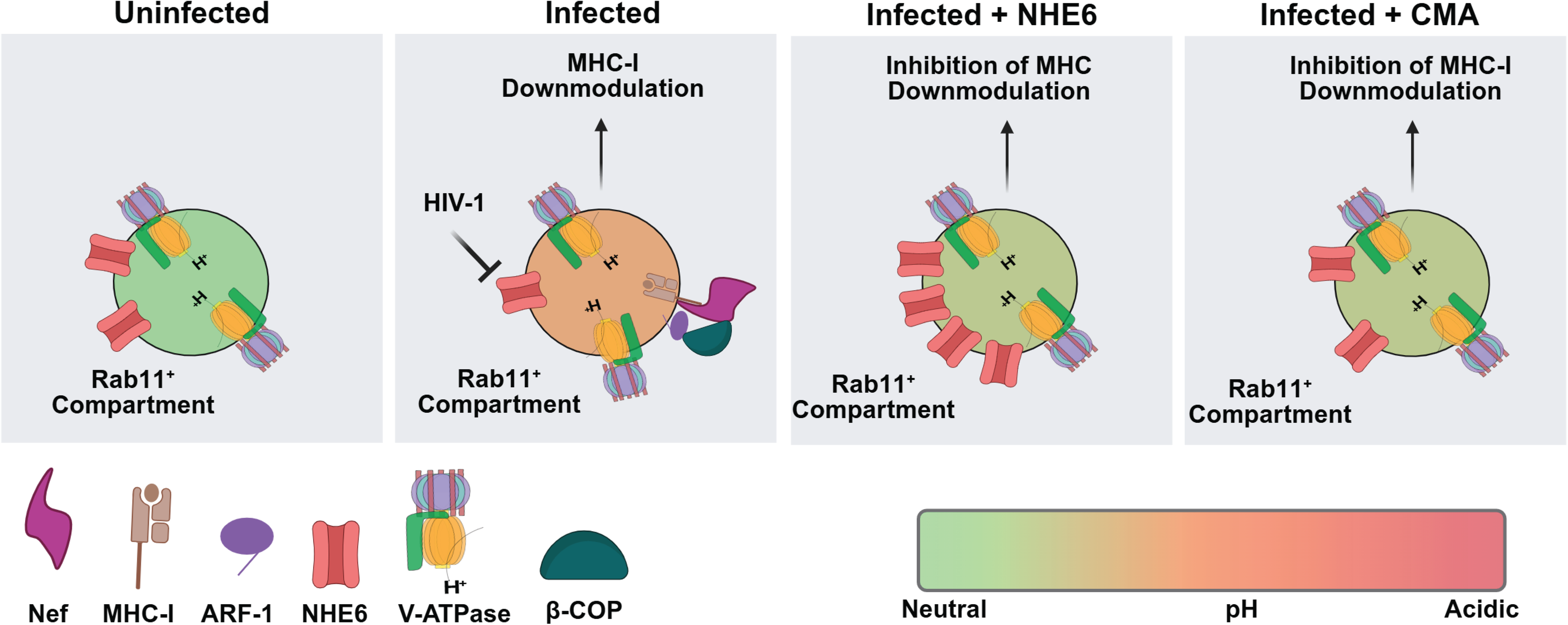
Model of the impact of neutralizing endosomal pH on Nef-dependent MHC-I Trafficking. Our findings suggest that HIV-1 infection triggers acidification of Rab11^+^ compartments by downmodulation of the 50 kDa form of NHE6. Neutralization of Rab11^+^ compartments via NHE6 overexpression reverses Nef-dependent MHC-I downmodulation by reducing recruitment of Nef to Rab11^+^ compartments and disrupting complex formation between Nef, β-COP, and ARF-1. Low-dose CMA similarly reduces acidity of Rab11^+^ compartments at concentrations that reverse Nef-dependent MHC-I trafficking.

We further found that selective neutralization of the mid and *trans*-Golgi network via overexpression of NHE8 also reduced Nef-dependent MHC-I downmodulation. This finding aligns with previous studies, as the role of the *trans-*Golgi network in Nef-mediated mis-trafficking of MHC-I is well established^24,26^. Similarly, CMA likely neutralizes compartments beyond those that were Rab11^+^ and occupied by NHE6, the primary focus of our study. While further elucidation of compartments targeted by CMA is needed, our emphasis on Rab11^+^ endosomal compartments occupied by NHE6 is justified for two reasons: (i) the role of the *trans-*Golgi network’s involvement in Nef-dependent MHC-I trafficking is already well characterized, whereas the role of the NHE6/Rab11^+^ compartment remains unclear, and (ii) HIV selectively targets the 50 kDa form of NHE6.

### The role of the NHE6/Rab11^+^ compartments in Nef-dependent MHC-I trafficking

Excitingly, we found that neutralization of Rab11^+^ compartments via NHE6 overexpression disrupted complex formation between β-COP, ARF-1, and Nef. Thus, our results highlight a critical function for this organelle in Nef-dependent MHC-I trafficking. This conclusion is further supported by data demonstrating that Nef localized to Rab11^+^ compartments in a pH-dependent manner, a process that was effectively disrupted by NHE6 overexpression.

### NHE6 expression is altered during infection by multiple viruses

Prior studies have reported that other viral infections impact NHE protein expression. Notably, proteomic analysis of cells infected with Enterovirus 71 revealed a significant reduction in NHE6 levels^27^. Similarly, decreased NHE6 protein expression was observed in the brains of mice infected with Chikungunya virus^28^, and diminished NHE6 gene expression was documented following Epstein-Barr virus infection^29^. While the full implications of diminished NHE6 expression in viral infections remain unclear, the consistent finding that multiple, diverse viral pathogens suppress NHE6 strongly suggests its important role in the host response to viral infection. Further investigation into the potential antiviral functions of NHE6 could uncover novel roles for this protein beyond its regulation of endosomal pH.

In contrast to NHE6, significant upregulation of *SLC9A9* expression, the gene encoding NHE9, has been observed in non-human primates challenged with several human-isolated strains of H5N1 avian flu viruses^30,31^. Likewise, NHE9 mRNA levels were markedly elevated in extracellular vesicles secreted from cells infected with West Nile virus^32^. These findings suggest that, beyond viruses directly targeting NHE expression, host cells may actively modulate NHE production in response to infection. Future exploration of the interplay between virus-induced alterations and cell-intrinsic regulation of NHE expression will be critical for unraveling the roles of these exchangers in viral pathogenesis and host defense.

### Nef and V-ATPase

It is noteworthy that Nef has been reported to interact with the H subunit of V-ATPase^33^, and these interactions may be relevant to endosomal pH regulation. Although this finding could suggest that Nef contributes to the acidification of certain endosomal compartments, we have thus far been unable to confirm the interaction between Nef and V-ATPase subunit H. Furthermore, we found no Nef-dependent difference in acidification of transferrin^+^ compartments.

While our study delineates a role for Rab11^+^ compartments in Nef-mediated trafficking of MHC-I to the lysosome, further research is needed to elucidate the functions of this and define other Nef-occupied compartments. Although we focused on the impact of reduced acidity in NHE6/Rab11^+^ compartments, it is likely that the pH of other organelles, such as the *trans*-Golgi network, is important as indicated by the effect of NHE8 overexpression on Nef activity. Additionally, late endosomes, and as-yet-unidentified compartments may also influence MHC-I trafficking by Nef. This is particularly important for understanding the mechanism of action of CMA in blocking Nef-dependent MHC-I downmodulation, as CMA likely impacts V-ATPase activity across multiple compartments. However, current tools to monitor compartment-specific pH changes remain limited.

Moreover, the precise mechanism by which HIV promotes NHE6 downmodulation, as well as the potential antiviral function of NHE6, warrants further investigation and represents an exciting avenue for future research. Although significant work remains to fully understand the interplay among HIV-1 Nef, endosomal compartment acidification, and virus-induced NHE6 downmodulation, our discovery provides novel insights into the critical role of NHE6/Rab11^+^ compartmental pH in Nef-dependent MHC-I trafficking.

## RESOURCE AVAILABILITY

### Lead contact

Further information and requests for resources and reagents should be directed to and will be fulfilled by the lead contact, Kathleen Collins, MD, PhD.

### Materials availability

All unique reagents generated in this study are available from the lead contact with a completed Materials Transfer Agreement.

### Data and code availability

- All data generated and reported in this paper are available from the lead contact upon request.
- This paper does not report original code.
- Any additional information required to reanalyze the data reported in this work paper is available from the lead contact upon request.

## Supporting information

Virus-induced vesicular acidification enhances HIV immune evasion._SI

## ACKNOWLEDGEMENTS

This research was funded by NIH grants R01 AI148383 to K.L.C, 1 TL1 DK 136046-1 to M.E.Y.M. G.G.F. was supported by the National Institute Of General Medical Sciences of the National Institutes of Health under Award Number T32GM149391. The content is solely the responsibility of the authors and does not necessarily represent the official views of the National Institutes of Health. The University of Michigan Flow Cytometry Core and the University of Michigan Microscopy Core provided access to instruments and technical support. Eurofins sequenced recombinant DNA constructs. We thank S.-J.-K. Yee (City of Hope National Medical Center) for providing pCMV-HIV-1. We thank Nancy Hopkins (Massachusetts Institute of Technology) for providing VSV-G. Figure 6A was created with BioRender.com. We would like to thank Matthew Huston for his help in characterizing the NHE overexpressing viruses in T cell lines. We would also like to thank Ricardo de Souza Cardoso for his expertise and help in quantifying the extent of colocalization between NHE6 and various organelle markers presented. Additionally, we want to thank Mark Painter for his ideas regarding the role of NHEs in Nef-dependent MHC-I downmodulation.

## AUTHOR CONTRIBUTIONS

M.E.Y.M. conducted experiments, analyzed data, and wrote the manuscript. G.G.F. conducted experiments, analyzed data, and assisted in writing the methods of the manuscript. G.E.Z. conducted experiments, analyzed data, and assisted in writing the methods section of manuscript. C.C. developed and initially tested the tagged Nef used in these studies. K.L.C. assisted in experimental design, data analysis and writing of the manuscript.

## FIGURE TITLES AND LEGENDS

**Supplemental Figure S1.** Characterization of NHE6 expression. Related to Figure 1. (A) General gating strategy for flow cytometry experiments. (B) NHE6 peptide epitope competitively blocks NHE6 antibody binding. Western blot analysis of the indicated amounts of CEM-A2 cell lysates incubated with primary NHE6 antibody that was preincubated with solvent or a peptide epitope of the NHE6 antibody at 10X excess of the antibody amount (100 µg of peptide) or 5X excess the antibody amount (50 µg of peptide). (C) NHE6 peptide epitope does not inhibit binding of non-NHE6 antibodies. Western blot analysis of the indicated amounts of CEM-A2 cell lysates incubated with primary NHE8 antibody that was preincubated with solvent or a peptide epitope of the NHE6 antibody as described in (B). (D) NHE6 co-localizes with transferrin. Representative confocal fluorescence microscopy images of untransduced CEM-A2 cells incubated with transferrin AF-647 for 15 minutes (time point of our assay) then stained for Rab5 and Rab11 (top panel). Confocal fluorescence microscopy images of CEM-A2 cells transduced with ΔGPE-IRES-NHE6 and incubated with transferrin AF-647 for 15 minutes, then stained for NHE6 (bottom). (E) Transferrin colocalizes with Rab5 and Rab11. Summary graph of colocalization of transferrin with Rab5 and Rab11. Colocalization was quantified as described in Figure 3F. A spots mask was assigned to transferrin, Rab5, and Rab11. The number of transferrin spots colocalized with Rab5 or Rab11 was divided by the total number of transferrin spots and graphed. Each point represents one image. Each symbol represents quantification of colocalization of transferrin with Rab5 or Rab11 from the same image. A total of 6 images were analyzed. The number of cells analyzed per imaged ranged from 2-4. 1 biological replicate was performed.

**Supplemental Figure S2.** Characterization of NHE6 and 8 expression. Related to Figure 2. (A) Full confocal microscopy images from Figure 2A, and C showing DAPI staining, bright field, and composite images. (B) Confirmation of NHE6 and NHE8 overexpression by western blot. Western blot analysis of lysates from CEM-A2 cells 48 hr post transduction with the indicated viruses and subjected to FACS analysis to isolate GFP+ (transduced) or GFP– (untransduced) cells and probed for the indicated NHEs. (C) NHE6 overexpression does not alter the pH of transferrin+ compartments in uninfected cells. Summary graph of transferrin+ compartment acidity of primary CD4+ T cells 3 days post transduction with the indicated viruses that either overexpress FLAG-tagged NHE6 (LeGo-BFP-NHE6-FLAG) or empty vector (LeGo-BFP). Cells were stained with pHrodo-transferrin and AF-647 transferrin. pHrodo transferrin MFIs for BFP+ (transduced) and BFP-(untransduced) cells were normalized to the corresponding AF-647 transferrin MFIs and graphed. One biological replicate was performed in technical triplicate. (D) NHE6 overexpression does not alter surface MHC-I expression in uninfected cells. Summary graph of surface MHC-I expression of primary CD4+ T cells 3 days post transduction with the indicated viruses. MHC-I MFIs for BFP+ (transduced) and BFP– (untransduced) cells were obtained and graphed. One biological replicate was performed in technical triplicate.

**Supplemental Figure S3.** Characterization of colocalization of NHE6 and NHE8 in CEM-A2 cells. Related to Figure 3. (A) HIV infection does not alter AP-1 staining. (Top panel) Full confocal images from Figure 3A including composite, bright field, and DAPI staining. (Bottom panel) NHE8 and AP-1 staining of mock transduced CEM-A2 cells. (B) Summary graph of quantification of colocalization between overexpressed NHE8 and AP-1 γ. Colocalization was quantified as described in Figure 3F. A spots mask was assigned for NHE8 and AP-1 γ staining. The number of NHE8 spots that colocalized with AP-1 γ was divided by the total number of NHE8 spots and graphed. Each point represents 1 image. A total of 12 images were analyzed. The number of cells quantified per image ranged from 1-4. 1 biological replicate was performed. (C) (Top panel) Full confocal images from Figure 3B including composite, bright field, and DAPI staining. (Bottom panel) staining of NHE6 and Rab11 in mock transduced CEM-A2 cells. (D) Full confocal images from Figure 3C including composite and DAPI staining. (E) Full confocal images from Figure 3D including composite and DAPI staining. (F) Full confocal images from Figure 3E including composite and DAPI staining.

**Supplemental Figure S4.** Overexpressed NHE9 does not colocalize with Rab7 in CEM-A2 cells. Related to Figure 3. (A) Representative confocal fluorescence microscopy images of CEM-A2 cells 48 hr post transduction with the indicated viruses and stained for NHE9 and Rab7. (B) Summary graph of quantification of expression of NHE8 48 hr post transduction of CEM-A2 cells with the indicated viruses. CTCF and fold change were calculated as described in Figure 2B. At least 30 cells were imaged. (C) Western blot analysis of CEM-A2 lysates 48 hr post transduction with the indicated viruses and sorted for infected (GFP+) cells. (D) Representative confocal fluorescence microscopy images of CEM-A2 cells 48 hr post transduction with the indicated virus then stained for NHE9 and Rab7. (E) Summary graph of quantification of colocalization between overexpressed NHE9 and Rab7. Colocalization was quantified as described in Figure 3F. A spots mask was assigned to NHE9 and Rab7 staining. The number of NHE9 spots colocalized with Rab7 was divided by the total number of NHE9 spots and graphed. Each point represents 1 image. 10 images were analyzed. The number of cells analyzed per image ranged from 1-3. 1 biological replicate was performed.

**Supplemental Figure S5.** NHE6 overexpression does not alter expression of β-COP or ARF-1. Related to Figure 5. (A) Western blot analysis of lysates from CEM-A2 cells 48 hr post transduction with the indicated viruses and subjected to FACS analysis to isolate the GFP+ (transduced) or GFP– (untransduced) populations. (B) Summary graph of β-COP protein expression from western blot analysis described in (A). β-COP signal was normalized to the corresponding vinculin signal and graphed. 3 biological replicates were performed. (C) Summary graph of ARF-1 protein expression from western blot analysis described in (A). ARF-1 signal was normalized to the corresponding vinculin signal and graphed. 3 biological replicates were performed. Statistical significance for (D) and (E) was determined with a One Way ANOVA mixed-effects analysis with Dunnett correction. * p < 0.05, ** p < 0.01, *** p < 0.001, **** p < 0.0001

**Supplemental Figure S6.** Characterization of colocalization between Nef and Rab11 with and without NHE6 overexpression. Related to Figure 6. (A) Full confocal images without (top panel) or with (bottom panel) pseudocoloring of Rab11 from Figure 6F (top panel) including composite and DAPI staining. (B) Full confocal images without (top panel) or with (bottom panel) pseudocoloring of Rab11 from Figure 6F (bottom panel) including composite and DAPI staining.

## STAR⍰Methods

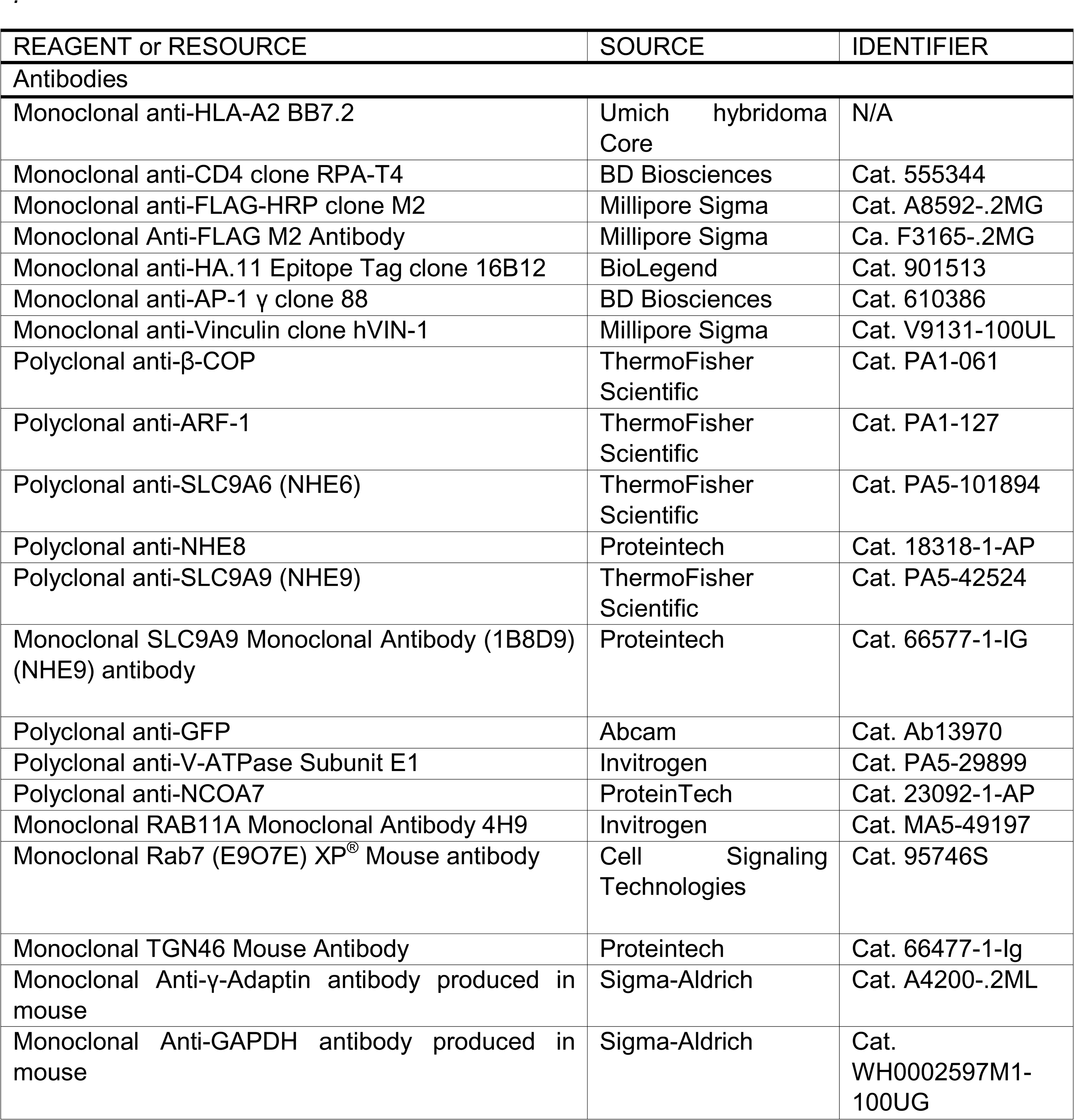
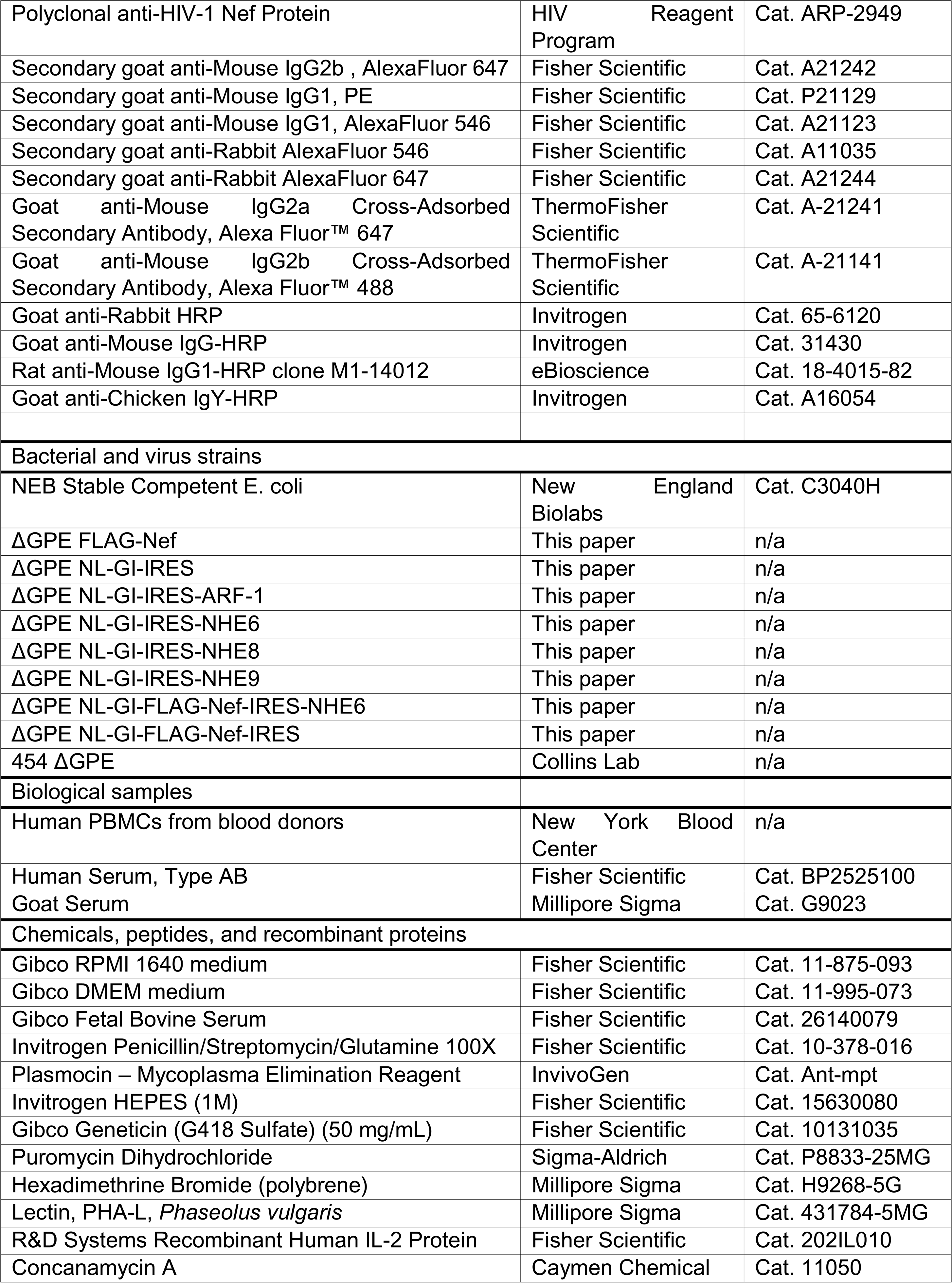
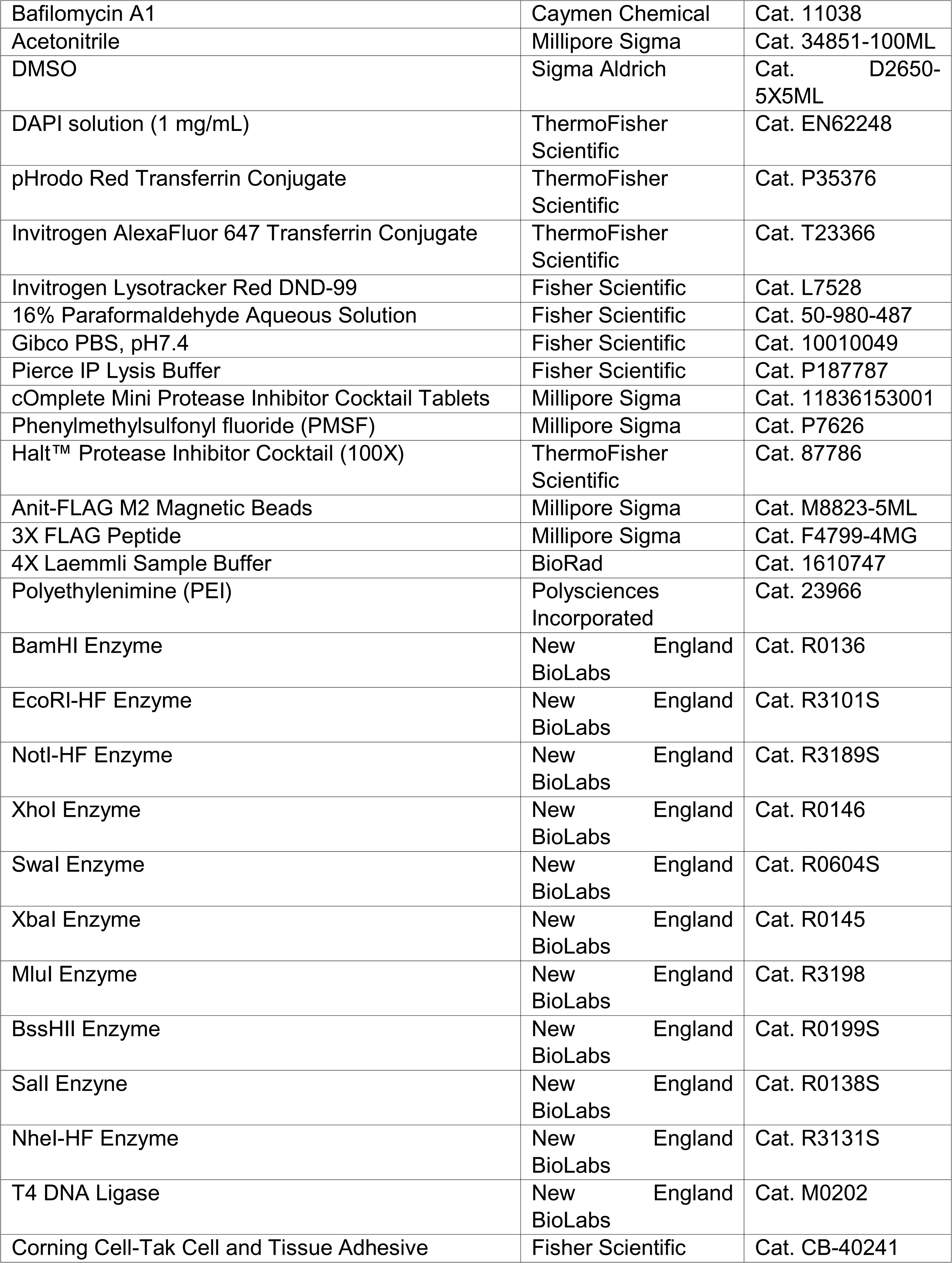
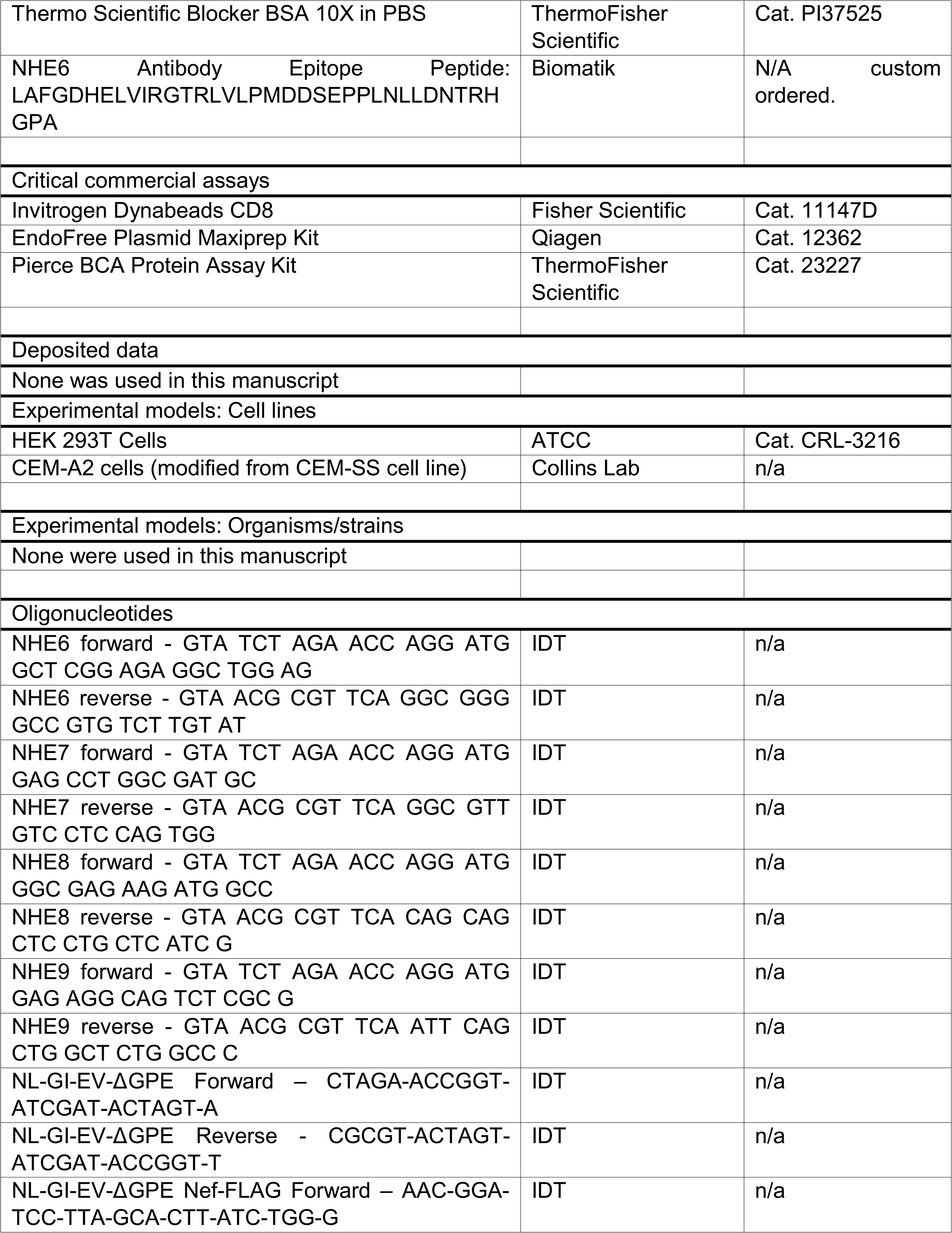
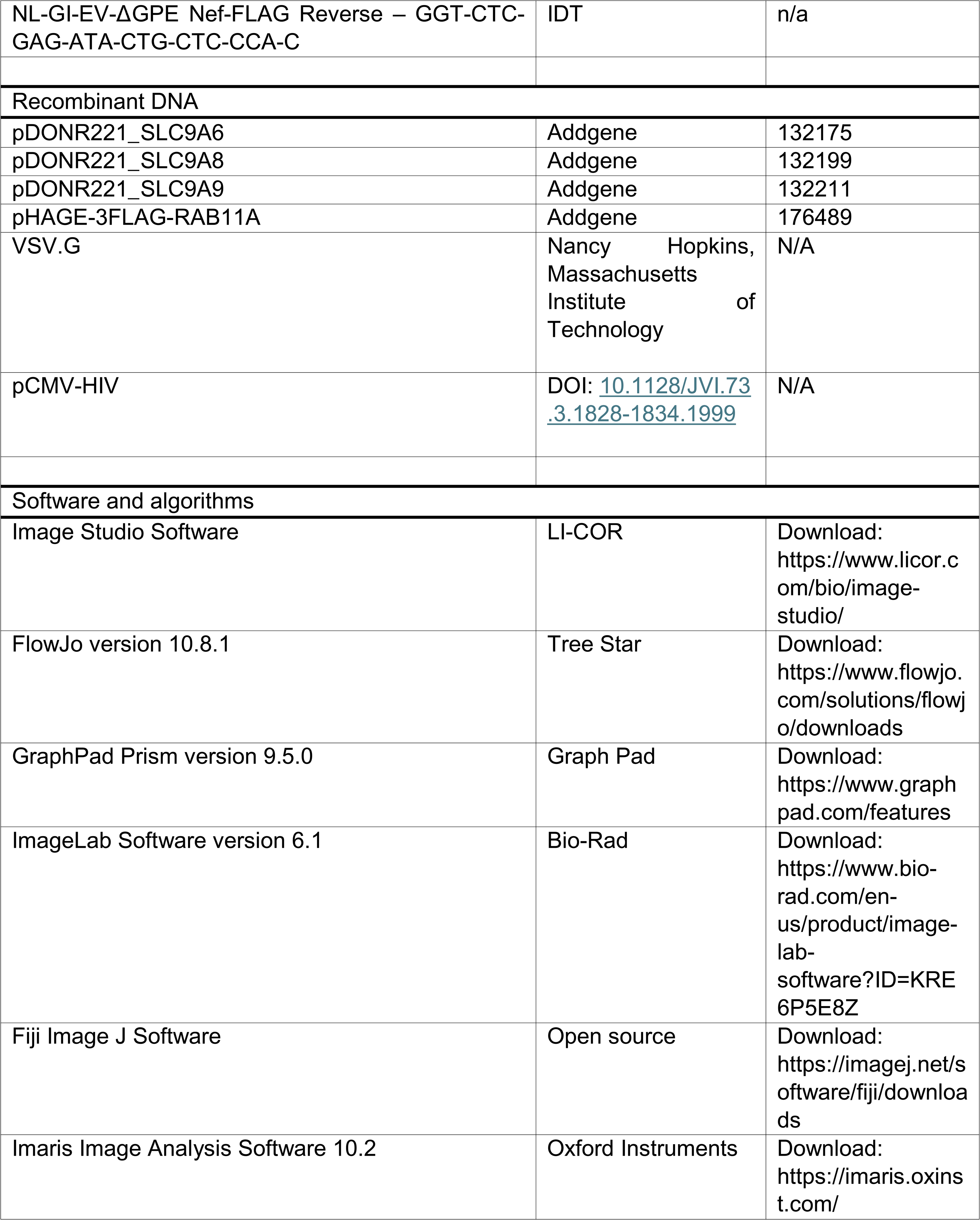
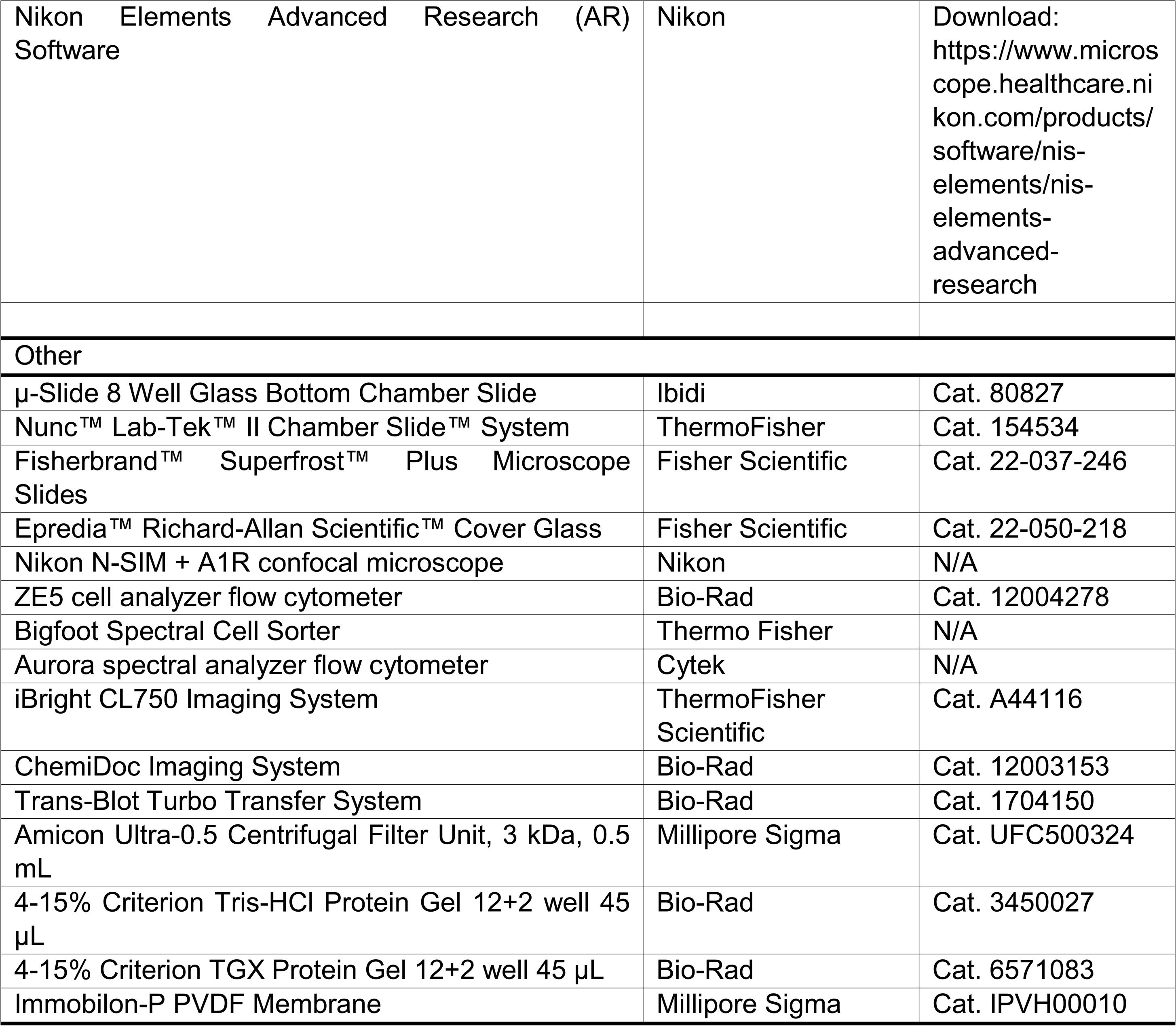
Key resources table.

### Experimental model and study participant details

#### Cell lines and Primary Cells

All cells were maintained at 37 °C in 5% CO_2_ humidified atmosphere. CEM cells engineered to express HA-tagged HLA-A2 (referred to as CEM-A2) were maintained in R10-Geneticin medium prepared as follows: RPMI 1640 medium (Fisher Scientific, USA) supplemented with Plasmocin (2.6 µg/mL, InvivoGen), 8.8 mM HEPES (Fisher Scientific), 0.87 U/mL Penicillin, 0.86 µg/mL streptomycin, 0.25 mg/mL L-glutamine (Penicillin-Streptomycin-Glutamine, Fisher Scientific), 10% fetal bovine serum (Fisher Scientific), and 0.95 mg/mL Geneticin (Fisher Scientific). Media was sterile filtered before use.

Frozen stocks of HLA-A2^+^ peripheral blood mononuclear cells (PBMCs) obtained from ficoll of Leukopaks from New York Blood Center were thawed and rested in R10 medium (prepared as described above, except without geneticin) overnight. The following day, cells were CD8 depleted by washing the cells once in with MACs buffer (2% FBS and 1 mM EDTA in PBS), adding 25 µL CD8 Dynabeads (Fisher Scientific) per 10^6^ PBMCs and washing twice with MACs buffer. PBMCs were resuspended at a density of 10^6^ cells/mL and rotated with the washed CD8 Dynabeads at 4 °C for 30 minutes. The supernatant containing only CD4^+^ cells was removed from the beads, pelleted, and resuspended in 20 mL of R10 supplemented with 5 µg/mL PHA (Millipore Sigma). The following day, 15 mL of media was removed from the flask and replaced with 5 mL of R10 supplemented with 100 U/mL IL-2 (final IL-2 concentration in the flask was 50 U/mL, Fisher Scientific). 72 hours post-PHA treatment, cells were used for downstream assays as described.

HEK 293T cells (ATCC) were maintained in D10, which was prepared like R10 except with DMEM medium (Fisher scientific) and without HEPES.

#### Generation of a Stable Cell Line Expressing FLAG-Tagged Rab11A

Virus was prepared with pHAGE-3FLAG-RAB11A as described below. CEM-A2 cells were transduced with the virus as described below. 48 hours post transduction, cells were resuspended at a density of 500,000 cells/mL in R10-Geneticin + 1 µg/mL puromycin (Sigma-Aldrich). Viability was monitored every day for 3 days and then cells were split with R10-Geneticin + 1 µg/mL puromycin at a density of 200,000 cells/mL. Cells were maintained in this media for 5 passages and were then transferred into R10-Geneticin + 0.5 µg/mL puromycin for long-term culture. Cells were maintained in this media.

### Method details

#### Cloning of ΔGPE-IRES-NHEs

NL4.3-GFP-IRES-ARF-1-ΔE virus^25^ was truncated to remove a section of the Gag-Pol gene by digesting with SwaI (NEB), gel purifying the truncated fragment and re-ligating to produce ΔGPE NL-GI-IRES-ARF-1. ARF-1 was then replaced with NHEs in ΔGPE NL-GI-IRES-ARF-1 using XbaI and MluI (NEB) cloning sites to obtain ΔGPE-IRES-NHE6/8/9. NHEs were obtained from Addgene (#132175, #132199, #132211). The following primers were used to amplify the NHE sequences and add the XbaI and MluI sites.

NHE6 forward – GTA TCT AGA ACC AGG ATG GCT CGG AGA GGC TGG AG

NHE6 reverse – GTA ACG CGT TCA GGC GGG GCC GTG TCT TGT AT

NHE8 forward – GTA TCT AGA ACC AGG ATG GGC GAG AAG ATG GCC

NHE8 reverse – GTA ACG CGT TCA CAG CAG CTC CTG CTC ATC G

NHE9 forward – GTA TCT AGA ACC AGG ATG GAG AGG CAG TCT CGC G

NHE9 reverse – GTA ACG CGT TCA ATT CAG CTG GCT CTG GCC C

ΔGPE NL-GI-IRES-ARF-1 and NHE inserts from PCR were double digested with XbaI and MluI. Following gel purification, digestion products were ligated according to the NEB ligation protocol with T4 DNA Ligase, transformed into NEB Stable Competent E. coli, and confirmed by Sanger sequencing and restriction digest.

#### Cloning of ΔGPE-IRES

Primers were designed to replace ARF-1 in ΔGPE NL-GI-IRES-ARF-1 with multiple cut sites (AgeI, ClaI, SpeI) rather than a protein of interest. The resulting construct is called ΔGPE NL-GI-IRES.

Forward – CTAGA-ACCGGT-ATCGAT-ACTAGT-A

Reverse – CGCGT-ACTAGT-ATCGAT-ACCGGT-T

Primers were annealed together and ligated with ΔGPE NL-GI-IRES-ARF-1 digested with XbaI and MluI then ligated after gel purification with T4 DNA ligase, following the NEB ligation protocol. The resulting product was transformed into NEB Stable Competent E. coli and confirmed by Sanger sequencing and restriction digest.

#### Cloning of ΔGPE-FLAG-Nef-IRES and ΔGPE-FLAG-Nef-IRES-NHE6

The following primers were used for PCR to amplify FLAG tagged Nef from a synthetic gBlock construct:

Forward – AAC-GGA-TCC-TTA-GCA-CTT-ATC-TGG-G

Reverse – GGT-CTC-GAG-ATA-CTG-CTC-CCA-C

gBlock sequence:

AACGGATCCTTAGCACTTATCTGGGACGATCTGCGGAGCCTGTGCCTCTTCAGCTACCACCGCTTGAGAGACTTACTCTTGATTGTAACGAGGATTGTGGAACTTCTGGGACGCAGGGGGT

GGGAAGCCCTCAAATATTGGTGGAATCTCCTACAGTATTGGAGTCAGGAACTAAAGAATAGTGCTGTTAACTTGCTCAATGCCACAGCCATAGCAGTAGCTGAGGGGACAGATAGGGTTATAGAAGTATTACAAGCAGCTTATAGAGCTATTCGCCACATACCTAGAAGAATAAGACAGGGCTTGGAAAGGATTTTGCTATAAGATGGGTGGCAAGTGGTCAAAAAGTAGTGTGATTGGATGGC CTGCTGTAAGGGAAAGAATGAGACGAGCTGAGCCAGCAGCAGATGGG**GACTACAAAGAC GATGACGACAAG**GTGGGAGCAGTATCTCGAGacc (Bolded letters indicate FLAG sequence. Letters highlighted in yellow represent the start codon for Nef.)

ΔGPE-IRES-NHE6, ΔGPE-IRES, and the PCR product were digested with XhoI and BamHI (NEB), gel extracted, and the products were ligated according to NEB ligation protocol with T4 DNA Ligase. The resulting product was transformed into NEB Stable Competent E. coli and confirmed by Sanger sequencing and restriction digest.

#### Cloning of ΔGPE FLAG-Nef

A FLAG tag was inserted into the N-terminal flexible loop of Nef with the following synthetic gBlock sequence:

GGATCCTTAGCACTTATCTGGGACGATCTGCGGAGCCTGTGCCTCTTCAGCTACCACCGCTTGAGAGACTTACTCTTGATTGTAACGAGGATTGTGGAACTTCTGGGACGCAGGGGGTGGGAAGCCCTCAAATATTGGTGGAATCTCCTACAGTATTGGAGTCAGGAACTAAAGAATAGTGCTGTTAACTTGCTCAATGCCACAGCCATAGCAGTAGCTGAGGGGACAGATAGGGTTATAGAAGTATTACAAGCAGCTTATAGAGCTATTCGCCACATACCTAGAAGAATAAGACAGGGCTTGGAAAGGATTTTGCTATAAGATGGGTGGCAAGTGGTCAAAAAGTAGTGTGATTGGATGGCCT GCTGTAAGGGAAAGAATGAGACGAGCTGAGCCAGCA**GACTACAAAGACGATGACGACAA G**GCAGATGGGGTGGGAGCAGTATCTCGAG (Bolded letters indicate FLAG sequence. Letters highlighted in yellow represent that start codon for Nef.)

The gblock and NL4.3 ΔGPE backbone were double digested with BamHI and XhoI (NEB). The digested products were then ligated with T4 DNA ligase following the NEB ligation protocol. The resulting product was transformed into NEB Stable competent E. coli and confirmed by Sanger sequencing and restriction digest.

#### Cloning of LeGo BFP NHE6-FLAG

C-terminal FLAG-tagged NHE6 was inserted into a LeGo-iBFP expression vector with the following synthetic gblock sequence:

TGGTGGTACGGGAATTCATGGCTCGGAGAGGCTGGAGGCGCGCCCCTCTGCGGAGAGGAGTGGGAAGCTCCCCTAGGGCAAGGCGCCTGATGAGACCACTGTGGCTGCTGCTGGCAGTGGGCGTGTTCGATTGGGCAGGAGCATCTGACGGAGGAGGAGGAGAGGCCCGGGCCATGGATGAGGAGATCGTGAGCGAGAAGCAGGCCGAGGAGTCCCACAGACAGGACTCTGCCAATCTGCTGATCTTCATCCTGCTGCTGACACTGACCATCCTGACCATCTGGCTGTTTAAGCACCGGAGAGCCAGGTTCCTGCACGAGACAGGCCTGGCCATGATCTACGGACTGCTGGTGGGACTGGTGCTGCGCTATGGCATCCACGTGCCATCCGATGTGAACAATGTGACCCTGTCTTGCGAGGTGCAGTCTAGCCCCACCACACTGCTGGTGACATTCGACCCTGAGGTGTTCTTTAATATCCTGCTGCCCCCTATCATCTTTTACGCCGGCTATTCCCTGAAGAGGCGCCACTTCTTTCGGAACCTGGGCTCTATCCTGGCCTACGCCTTTCTGGGCACCGCCATCAGCTGCTTCGTGATCGGCTCCATCATGTATGGCTGCGTGACCCTGATGAAGGTGACCGGCCAGCTGGCCGGCGATTTCTACTTTACCGACTGTCTGCTGTTCGGCGCCATCGTGTCCGCCACAGATCCAGTGACCGTGCTGGCCATCTTCCACGAGCTGCAGGTGGACGTGGAGCTGTATGCCCTGCTGTTTGGCGAGAGCGTGCTGAATGACGCCGTGGCCATCGTGCTGTCCTCTAGCATCGTGGCATACCAGCCAGCAGGCGATAACTCTCACACATTTGACGTGACCGCCATGTTCAAGAGCATCGGCATCTTCCTGGGCATCTTTTCTGGCAGCTTCGCCATGGGAGCAGCAACAGGAGTGGTGACCGCCCTGGTGACAAAGTTTACCAAGCTGAGGGAGTTCCAGCTGCTGGAGACAGGCCTGTTCTTTCTGATGTCCTGGTCTACCTTTCTGCTGGCAGAGGCATGGGGCTTCACAGGAGTGGTGGCCGTGCTGTTTTGCGGCATCACACAGGCCCACTACACCTATAACAATCTGAGCACAGAGTCCCAGCACCGCACCAAGCAGCTGTTTGAGCTGCTGAACTTCCTGGCCGAGAATTTCATCTTTTCCTATATGGGCCTGACACTGTTCACCTTTCAGAACCACGTGTTTAATCCTACCTTCGTGGTGGGCGCCTTTGTGGCCATCTTCCTGGGCAGGGCCGCCAACATCTACCCACTGAGCCTGCTGCTGAATCTGGGCCGGAGATCTAAGATCGGCAGCAATTTTCAGCACATGATGATGTTCGCAGGACTGAGGGGAGCAATGGCCTTCGCCCTGGCCATCAGGGATACAGCCACCTACGCCCGCCAGATGATGTTCTCCACCACACTGCTGATCGTGTTCTTTACCGTGTGGGTGTTTGGAGGAGGAACCACAGCAATGCTGAGCTGCCTGCACATCCGGGTGGGCGTGGATTCCGACCAGGAGCACCTGGGAGTGCCAGAGAACGAGAGGAGGACCACAAAGGCAGAGTCCGCCTGGCTGTTTAGAATGTGGTACAACTTCGACCACAATTATCTGAAGCCACTGCTGACCCACTCTGGACCACCACTGACCACAACCCTGCCAGCATGCTGTGGACCTATCGCAAGATGTCTGACCAGCCCTCAGGCCTATGAGAACCAGGAGCAGCTGAAGGACGATGACTCCGATCTGATCCTGAATGATGGCGACATCTCTCTGACCTACGGCGACAGCACAGTGAACACCGAGCCAGCCACATCCTCTGCCCCACGGAGATTCATGGGCAACAGCTCCGAGGATGCACTGGACAGGGAGCTGGCCTTTGGCGATCACGAGCTGGTCATCAGGGGAACCAGGCTGGTGCTGCCAATGGATGACAGC GAGCCTCCACTGAACCTGCTGGACAATACAAGACACGGCCCCGCC**GACTACAAAGACGA TGACGACAAG**TGAGCGGCCGCTACGTAAATTCC (Bolded letters indicated FLAG sequence).

The gblock and LeGo-iBFP backbone were double digested with EcoRI and NotI (NEB). The digested products were then ligated following gel purification with T4 DNA ligase following the NEB ligation protocol. The resulting product was transformed into NEB Stable competent E. coli and confirmed by Sanger sequencing and restriction digest.

#### Virus Production

2 million HEK 293T cells were plated on a 10 cm dish and adhered overnight. The following day, 4 µg of VSV-G plasmid, 4 µg of pCMV-HIV plasmid, and 4 µg of the viral genome of interest were diluted in 952 µL of 150 mM NaCl. 48 µL of a 1 mg/mL stock of polyethylenimine (PEI, Polyscience Incorporated) was added to the solution and this was gently vortexed. The transfection solution incubated at room temperature for 15 minutes and 1 mL was added to the plate of cells. The transfection solution was scaled accordingly to facilitate transfection of multiple plates at once. Cells incubated with the transfection solution for 16 hours. The media was then aspirated and replaced with fresh D10. 48 hours post-transfection, media containing the virus was pelleted to remove dead cells and aliquots were frozen at –80 °C until use. All viruses were titrated with CEM-A2 cells to determine volumes of virus required for desired infection rates.

#### Viral Transductions

CEM-A2 cells were resuspended in virus diluted in D10 to achieve the desired infection rate at a density of 1 million cells/mL. 1 mL or 500 µL of cells was added to the wells of 24 or 48-well plate (0.5-1 million cells/well). Cells centrifuged at 2,500 rpm for 2 hours at room temperature. Media was then aspirated and replaced with 1-2 mL of R10. Cells were cultured for 48-72 hours depending on the downstream application.

Primary CD4^+^ T cells were transduced as described above in a 24-well plate, except 4 µg/mL of polybrene (Millipore Sigma) was added to the virus diluted in D10. After the centrifugation step, media was aspirated and replaced with 1 mL of R10 supplemented with 50 U/mL of IL-2 (Fisher Scientific).

#### Fluorescence Confocal Microscopy

A 0.02 µg/µL stock of Cell-Tak (Fisher Scientific) was prepared by diluting a stock solution in sterile 0.1 M sodium bicarbonate buffer, pH 8.0. 3.9 µg of Cell-tak was added to the wells of an 8-chamber glass coverslip (Ibidi or ThermoFisher) or 20 µg was added to 22X22 glass coverslips (Fisher Scientific) and incubated at room temperature for 20 minutes. The solution was aspirated, and wells or coverslips were washed once with sterile water. The coverslips airdried at room temperature and when necessary, were wrapped in parafilm and stored overnight at 4 °C.

##### Antibodies used for Confocal Microscopy

Primary antibodies were diluted in 2% BSA (paraformaldehyde fixation described below) or 1% BSA (ethanol fixation described below). We utilized antibodies directed toward NHE6 (ThermoFisher Scientific diluted 1:100), NHE8 (ProteinTech, diluted 1:50), NHE9 (Thermofisher Scientific, diluted 1:50 or ProteinTech diluted 1:100), FLAG (for FLAG-tagged Nef, Sigma Millipore, diluted 1:400), Rab11A (Invitrogen, diluted 1:100), AP-1 γ (Sigma Aldrich, diluted 1:200), TGN46 (Proteintech diluted 1:200), and Rab7 (Cell Signaling Technologies, diluted 1:100). Secondary antibodies were diluted 1:250 in 2% BSA (for paraformaldehyde fixation described below) or 1:200 in 1% BSA (for ethanol fixation described below). For detection of NHE6, NHE8, and NHE9 (ThermoFisher Scientific) we utilized goat anti-rabbit AF-647 (Fisher Scientific) or goat anti-rabbit AF-546 (Fisher Scientific) secondary antibodies. For detection of NHE9 (ProteinTech) we utilized a goat anti-mouse IgG2a AF-647 (ThermoFisher Scientific) secondary antibody. For detection of Rab11A and AP-1 γ we utilized goat anti-mouse IgG2b AF-647 (ThermoFisher Scientific), goat anti-mouse IgG2b AF-488 (ThermoFisher Scientific), or goat anti-mouse IgG2b AF-546 (Fisher Scientific). For detection of Rab7, TGN46, and FLAG-tagged Nef we utilized goat anti-mouse IgG1 AF-546 (Fisher Scientific).

##### Transferrin uptake

Cells plated in a round bottom 96-well plate were washed once with Live Cell Imaging Buffer (LCIB, 140 mM NaCl, 2.5 mM KCl, 1.8 mM CaCl_2_, 1.0 mM MgCl_2_, 20 mM HEPES, pH 7.4 supplemented with 1% BSA right before use). Transferrin conjugated to AlexaFluor 647 (ThermoFisher Scientific) was diluted 1:200 in LCIB and incubated with the cells at 37 °C for 15 minutes. Cells were pelleted and washed once with LCIB. Cells were then subjected to paraformaldehyde fixation and antibody staining described below.

##### Paraformaldehyde fixation (NHE6 and Rab11, Rab7, or TGN46 and Nef and Rab11 staining)

Cells were fixed by adding 2% paraformaldehyde in PBS (PFA, Fisher Scientific) and incubating at room temperature for 20 minutes. Cells were permeabilized with 0.2% tween20 in PBS for 20 minutes at 37 °C, pelleted and washed once with 2% BSA in PBS. For the antibody staining step, cells were first pre-incubated on ice for 30 minutes with 2% BSA in PBS. Then, antibodies were diluted in 2% BSA in PBS as described above, added to the cells and were incubated for 20 minutes on ice. Cells were then pelleted and washed once with 2% BSA. Then, secondary antibodies were diluted in 2% BSA as described above, added to the cells, and were incubated on ice for 20 minutes, protected from light. Cells were then pelleted and washed once with 2% BSA. Nuclei were labeled by diluting a 1 mg/mL DAPI stock (ThermoFisher Scientific) 1:1,000 in PBS and incubating with the cells for 5 minutes at room temperature. Cells were washed once with 2% BSA, then resuspended in 200 µL 2% BSA and added to the wells of the cell-Tak coated coverslip. Cells were adhered to the coverslip for 30 minutes at room temperature. Cells stained with NHE6 and Rab11 were imaged immediately after adhering. For cells stained with NHE6 and Rab5, Rab7 or TGN46, the chamber was removed from the coverslip and a drop of ProlongGold^TM^ antifade (Fisher Scientific) was added to the samples and a coverslip was placed on top. The ProlongGold^TM^ antifade cured for 24 hours at room temperature and then cells were imaged. For cells stained with Rab11 and FLAG (Nef), cells adhered on a glass coverslip as described above, then a drop of ProlongGold^TM^ antifade was placed on a glass slide. The coverslip was mounted on the slide and the ProlongGold^TM^ cured for 24 hours at room temperature and cells were imaged.

##### Ethanol fixation (NHE8 and NHE9 staining and colocalization of NHE8 with AP-1 or NHE9 with Rab7)

Cells were resuspended in PBS at a density of 1,750 cells/µL and 200 µL was added to the wells of a cell-Tak coated coverslip (350,000 cells/well). Cells adhered to the coverslip for 20 minutes at 37 °C and were fixed by adding ice-cold 100% ethanol to the wells and incubating at room temperature for 10 minutes. Ethanol was aspirated and cells were washed 3 times with PBS. Cells were permeabilized by adding 0.2% tween20 in PBS and incubating for 5 minutes at room temperature. Cells were washed 3 times with PBS. Prior to antibody staining, cells were preincubated with 5% goat serum (Millipore Sigma) in PBS for detecting overexpression of NHE8 and NHE9 or 2% BSA to monitor colocalization of NHE8 with AP-1 or NHE9 with Rab7 for 1 hour at room temperature. Then, primary antibodies described above were diluted in 1% BSA in PBS and incubated with the cells for 2.5 hours at room temperature with gentle rocking. The antibodies were then aspirated, and cells were washed three times with PBS with rocking for 5 minutes at room temperature. Secondaries were diluted 1:200 in 1% BSA as described above and incubated with the cells for 1.5 hours at room temperature with gentle rocking, protected from light. The secondary was aspirated, and cells were washed three times with PBS with rocking for 5 minutes at room temperature. Nuclei were stained with DAPI as described above. Cells were washed once with PBS and 200 µL of PBS was added to each well.

##### Imaging

Slides were imaged with a Nikon N-SIM + A1R confocal microscope. Identical laser and gain settings were used across all images for each replicate of the experiment.

#### Flow Cytometry Staining

Nef-dependent MHC-I and/or CD4 downmodulation from the cell surface in response to all viral transductions was evaluated by flow cytometric analysis of transduced primary T cells. Cells were stained in a round bottom 96-well plate 72 hours post transduction. Surface expression of MHC-I and/or CD4 was monitored using antibodies directed against HLA-A2 (BB7.2, Umich Hybridoma Core, 1:2,000 dilution) and CD4 (BD Biosciences 1:500 dilution). All antibodies were diluted in FACS buffer (2% FBS, 1% human serum, 10 mM HEPES, and 0.025% sodium azide in PBS). The antibody cocktail was added to the cells and incubated on ice for 20 minutes. Cells were pelleted and washed once with FACS buffer prior to incubation with secondary antibodies [goat anti-mouse IgG2b AF-647 (to detect BB7.2, Fisher Scientific) and goat anti-mouse IgG1 PE (to detect CD4, Fisher scientific)] diluted 1:2,000 in FACS buffer. DAPI was added as a viability dye to the secondary antibody cocktail at a final concentration of 0.04 µg/mL. Cells were incubated with the cocktail for 20 minutes on ice, protected from light. Cells were washed once with FACS buffer and fixed by adding 2% PFA to the wells and incubating for 20 minutes at room temperature, protected from light. Cells were resuspended in FACS buffer and analyzed on a ZE5 cell analyzer (Bio-Rad) or an Aurora spectral analyzer (Cytek). A minimum of 10,000 live events were collected for each sample. All laser and gain settings were identical for all samples for each individual replicate of the experiment.

Changes in primary CD4^+^ T cell transferrin^+^ compartment pH in response to viral transduction were monitored 72 hours post-transduction with the indicated viruses. Cells were washed once with LCIB. Transferrin conjugated to pHrodo Red or AF-647 (ThermoFisher Scientific) was diluted 1:200 in LCIB and added to the wells. Cells were incubated for 15 minutes at 37 °C, pelleted and washed once with LCIB. Cells were resuspended in LCIB and analyzed without fixing on a ZE5 cell analyzer (Bio-Rad) or an Aurora spectral analyzer (Cytek). Laser and gain settings were identical for all samples for each individual replicate of the experiment.

To monitor neutralization of transferrin^+^ compartments in response to CMA or Bafilomycin A1 treatment, primary CD4^+^ T cells transduced with the indicated viruses and dosed with CMA (described below) or Bafilomycin A1 (described below) were pelleted in a round bottom 96-well plate 24 hours post compound treatment and stained exactly as described above.

To monitor neutralization of the lysosome in response to CMA or Bafilomycin A1 treatment or viral transduction, transduced primary CD4^+^ T cells treated with or without the compounds were collected for staining 72 hours post transduction in a 96-well round bottom plate. Lysotracker Red (Fisher Scientific) diluted 1:5,000 in PBS was added and cells were incubated for 1 hour at 37 °C. Cells were then pelleted and washed once with FACS buffer. DAPI diluted to a final concentration of 0.04 µg/mL in FACS buffer was added to the cells and incubated 5 minutes at room temperature, protected from light. Cells were pelleted and washed once with FACS buffer. Cells were fixed by adding 2% PFA in PBS and incubating at room temperature for 20 minutes. Cells were resuspended in FACS buffer and analyzed with a ZE5 cell analyzer (Bio-Rad) or an Aurora spectral analyzer (Cytek). Laser and gain settings were identical for all samples for each individual replicate of the experiment.

#### CMA Titrations

Primary HLA-A2^+^ CD4^+^ T cells were isolated as described above. 72 hours post PHA treatment, cells were transduced with ΔGPE FLAG-Nef virus as described in the viral transduction section. 2 days post transduction, cells were resuspended at a density of 2 million cells/mL in R10 + 100 U/mL IL-2. 50 µL of the cell suspension was added to the wells of a flat-bottom 96-well plate (100,000 cells/well). Two titrations of CMA were performed. For the first titration, CMA (Caymen chemical) was diluted in R10 to produce a 1.5 nM stock and acetonitrile (Millipore Sigma) was diluted the same way to produce a solvent control. CMA and acetonitrile were serial diluted 1:1 in R10 to produce concentrations ranging from 0.75 – 0.0029 nM. In an effort to obtain more data points near the IC_50_ for transferrin conjugated to pHrodo Red signal (indicative of transferrin^+^ compartment neutralization) a second titration of CMA and acetonitrile was prepared in which a 0.67 nM CMA stock was prepared in R10. This stock or acetonitrile was serial diluted 1:1.5 in R10 to produce a range of CMA concentrations from 0.33 – 0.017 nM. 50 µL of each CMA or acetonitrile concentration was added to the cells. Final media volume was 100 µL/well with 100,000 cells/well. For the first CMA titration, the final range of CMA concentrations was 0.38 nM – 0.001 nM. The final range of concentrations for the second CMA titration was 0.33 nM – 0.009 nM. The final IL-2 concentration was 50 U/mL. Cells were incubated with CMA for 24 hours and then stained for flow cytometric analysis of MHC-I expression and transferrin^+^ compartment neutralization as described above.

Since neutralization of the lysosome occurs at substantially higher concentrations of CMA than required to reverse Nef-dependent MHC-I downmodulation and neutralize transferrin^+^ compartments, a separate CMA titration was performed starting a higher concentration. Primary CD4^+^ were transduced with ΔGPE FLAG-Nef and added to the flat-bottom 96-well plate as descried above. CMA (Caymen Chemical) was diluted to produce a 20 nM stock in R10 and acetonitrile was diluted the same way to produce a solvent control. This stock and the acetonitrile were serial diluted 1:1 to produce concentrations of CMA ranging from 20 – 0.01 nM. CMA or acetonitrile at each concentration was added to the cells. The final volume in the wells was 100 µL with 100,000 cells/well. The final range of CMA concentrations was 10 – 0.005 nM. The final IL-2 concentration in the wells was 50 U/mL. Cells were incubated with CMA for 24 hours prior to staining with lysotracker as described above.

#### Bafilomycin A1 Titrations

Primary HLA-A2^+^ CD4^+^ T cells were isolated as described above. 72 hours post PHA treatment, cells were transduced with ΔGPE FLAG-Nef virus as described in the viral transduction section. 2 days post transduction, cells were resuspended at a density of 2 million cells/mL in R10 + 100 U IL-2/mL. 50 µL of the cell suspension was added to the wells of a flat-bottom 96-well plate (100,000 cells/well). Two titrations of Bafilomycin A1 (Baf A1) were performed. For the first titration, Baf A1 (Caymen chemical) was diluted in R10 to produce a 200 nM stock and DMSO (Sigma Aldrich) was diluted the same way as a solvent control. Baf A1 and DMSO were serial diluted 1:1 in R10 to produce concentrations ranging from 200 – 0.1 nM. In an effort to obtain more data points near the IC_50_ for transferrin conjugated to pHrodo Red signal (indicative of transferrin^+^ compartment neutralization) a second titration of Baf A1 and DMSO was prepared in which a 25 nM stock was prepared in R10. This stock or DMSO was serial diluted 1:1.5 in R10 to produce a range of Baf A1 concentrations from 25 – 0.6 nM. 50 µL of each Baf A1 or DMSO concentration was added to the cells. Final media volume was 100 µL/well with 100,000 cells/well. For the first Baf A1 titration, the final range of Baf A1 concentrations was 100 nM – 0.05 nM. The final range of concentrations for the second Baf A1 titration was 12.5 nM – 0.3 nM. The final IL-2 concentration was 50 U/mL. Cells incubated with Baf A1 for 24 hours and were then stained for flow cytometric analysis of MHC-I downmodulation, transferrin^+^ compartment neutralization, and lysotracker as described above.

#### Fluorescence-Activated Cell Sorting (FACS)

For all sorting experiments, cells were collected 48 hours post transduction and resuspended at a density of 5,000,000 cell/mL in MACs buffer. Cells were sorted based on GFP expression with a Bigfoot Spectral Cell Sorter (ThermoFisher Scientific) into tubes containing R10.

#### Western Blot Analysis

##### Lysate preparation, SDS PAGE and transfer

Cells were washed once with PBS, counted and lysed by adding 30 µL Pierce IP lysis buffer (ThermoFisher Scientific) with HALT protease inhibitor cocktail (ThermoFisher Scientific) and rocking on ice for 5 minutes. The lysate was clarified by centrifuging at 14,000 xg for 10 minutes at 4 °C. Lysates were mixed with 4X Laemmli Sample Buffer (Bio-Rad) and heated at 95 °C for 5 minutes prior to running on a 4-15% TGX criterion gel at 150 V for 1 hour. Protein was transferred to a PVDF membrane with a Trans-Blot Turbo Transfer system (Bio-Rad) using the StandardSD (1 Amp, 25 V, 30 min) method. The membrane was blocked with 5% fat free milk in TBST for 1 hour at room temperature.

##### Antibodies used for western blot analysis

All primary antibodies were diluted in 5% fat free milk in TBST and incubated overnight at 4 °C with gentle rocking. We utilized antibodies directed against NCOA7 (ProteinTech, 1:1,000). NHE6 (ThermoFisher, 1:100-1:500), NHE8 (ProteinTech 1;100-1:500), NHE9 (ThermoFisher, 1:100-1:500), V-ATPase subunit E1 (ThermoFisher, 1:500), Vinculin (Millipore Sigma, 1:1,000), GFP (Abcam, 1:1,000), Nef (AIDS research and reference reagent program, 1:100-1:500), β-COP (ThermoFisher Scientific, 1:100-1:1,000), ARF-1 (ThermoFisher Scientific,1:100 –1:1000), AP-1 γ (BD Biosciences, 1:100), HA.11 (BioLegend, 1:100), FLAG-HRP (Millipore Sigma, 1:1,000-1:10,000), and GAPDH (Sigma Aldrich, 1:1,000).

##### Membrane staining

The following day, the blots were washed 5X with TBST with rocking for 5 minutes per wash at room temperature. All secondaries conjugated to horseradish peroxidase were diluted 1:10,000 in 5% fat free milk in TBST and incubated with the blots for 1 hour at room temperature. For detecting NCOA7, NHE6, NHE9, V-ATPase subunit E1, Nef, β-COP, and ARF-1, blots were incubated with goat anti-rabbit IgG-HRP secondary (ThermoFisher Scientific). For detecting Vinculin, GAPDH, and AP-1 γ, blots were incubated with goat anti-mouse IgG-HRP secondary (ThermoFisher Scientific or R&D Systems respectively). For detecting GFP, blots were incubated with goat anti-Chicken IgY-HRP (ThermoFisher Scientific). For detecting HA.11, rat anti-mouse IgG1 (eBioscience) was used. Blots were washed 5X with TBST for 5 minutes per wash, incubated with ECL substrate and chemiluminescent signal was visualized with a Chemidoc Imaging system (Bio-Rad).

#### Co-Immunoprecipitation of Proteins Bound to Nef

25 million CEM-A2 cells were transduced with ΔGPE-FLAG-Nef-IRES-NHE6, ΔGPE-FLAG-Nef-IRES or mock transduced with D10 containing no virus as described above. To inhibit excessive degradation of MHC-I and other proteins of interest by the lysosome, 48 hours post transduction, cells were collected, counted, and resuspended in R10 supplemented with 35 mM NH_4_Cl at a density of 1 million cells/mL. 24 hours post NH_4_Cl treatment, equal number of cells for each condition were pelleted, washed with PBS, and transferred to a pre-weighed tube. PBS was aspirated and the weight of the cell pellet was calculated. The weight of the cell pellet (in mg) was then multiplied by 10 to calculate the appropriate volume of lysis buffer (i.e. a 50 mg pellet would be lysed in 500 µL of buffer). A subset of cells was subjected to flow cytometric analysis to check that transduction rates between the viruses were similar and Nef-dependent MHC-I downmodulation occurred as expected.

Peirce IP lysis buffer (ThermoFisher Scientific) was supplemented with 1 mM phenylmethylsulfonyl fluoride (PMSF, Millipore Sigma) and 1X protease inhibitor tablet (10X solution prepared by dissolving 1 tablet in 1 mL of Pierce IP buffer, Millipore Sigma). Cell pellets were resuspended in lysis buffer according to their weight as described above and rocked on ice for 5 minutes. Lysate was clarified by centrifuging at 14,000 xg for 10 minutes at 4 °C. A fraction of lysate was saved to run as an input control. 40 µL of magnetic FLAG resin (Millipore Sigma) per 25 million cells was washed three times with 4 resin volumes of lysis buffer. Lysates were added to the resin and rotated at 4 °C for 1.5 hours. Resin was separated from the lysate with a magnetic stand, lysate was aspirated, and resin was washed 3 times with 500 µL of modified IP buffer (Pierce IP buffer except with 0.1% NP-40 and 0.5% glycerol) with rotation at 4°C for 5 minutes. The resin was transferred to a clean tube for the final wash.

Protein was eluted from the resin by adding a 200 ng/µL stock of 3X FLAG peptide (Millipore Sigma) diluted in modified IP buffer and agitated at room temperature for 30 minutes. Resin was separated from the eluate with a magnetic stand and eluate was collected. Eluate was concentrated from 100 µL to 30 µL by centrifuging in Amicon centrifugal filter units with a 3 kDa molecular weight cutoff (Millipore Sigma) for 1 hour at 14,000 xG at 4 °C.

4X Laemmli Sample Buffer (Bio-Rad) was mixed with all the samples and heated at 95 °C for 5 minutes. Samples were run on a 4-15% gradient acrylamide gel (Bio-Rad) at 130 V for 90 minutes. Protein was transferred to PVDF membrane at 350 mAmp for 90 minutes. The membrane was then blocked with 5% fat free milk in TBST for 1 hour at room temperature and stained as described above.

#### Isolation of Intact Rab11^+^ Compartments

48 hours post transduction with ΔGPE-IRES or ΔGPE-IRES-NHE6, CEM-A2 cells stably expressing FLAG-tagged Rab11 (CEM-A2-FLAG-Rab11) were pelleted (∼17 million cells per sample). As a control, mock transduced CEM-A2-FLAG-Rab11 and CEM-A2 cells were included. Cells were washed 1X with KPBS (25 mM KCl, 100 mM potassium phosphate, pH 7.2) and were resuspended at a density of 1.2 million cells/mL in KPBS + HALT protease inhibitor (ThermoFisher Scientific) diluted to 1X final concentration and lysed with 50 strokes of a 2 mL Dounce homogenizer on ice. Lysate was clarified twice at 1,000 xG at 4 °C for 5 min. Aliquots of 60 µL of Anti-FLAG M2 Magnetic Beads (Millipore Sigma) were washed three times with 1 mL of lysis buffer. 20 µL of clarified lysate was saved to run as an input fraction and mixed with 10 µL of lysis buffer to produce a final volume of 30 µL. The remaining lysate was added to the beads and incubated for 50 minutes at 4 °C with rotation. Beads were separated from the lysate with a magnetic stand and unbound lysate was aspirated. Beads were washed three times with 500 µL of lysis buffer. Each wash rotated for 5 minutes at 4 °C and beads were transferred to a clean tube for the final wash. Protein was eluted from the beads by adding 30 µL of a 500 ng/µL stock of 3X FLAG peptide (Millipore Sigma) diluted in KPBS and agitated at room temperature for 45 minutes. Beads were separated from the eluate with a magnetic stand and the eluate fraction was collected. This procedure was taken and modified from dx.doi.org/10.17504/protocols.io.ewov14pjyvr2/v2. Samples were separated by SDS PAGE, transferred using a Trans-Blot Turbo Transfer system (Bio-RAD) and analyzed via western blot as described above.

#### NHE6 Peptide Competition of Antibody Binding

Because western blot analysis using the NHE6 antibody (ThermoFisher Scientific) produced numerous bands, we performed a peptide competition to identify bona fide NHE6 bands. A peptide corresponding to the epitope recognized by the antibody was custom order from Biomatik (sequence of peptide: LAFGDHELVIRGTRLVLPMDDSEPPLNLLDNTRHGPA). The peptide was dissolved in a 4:1 ratio of acetonitrile:water (solvent). 5 million CEM-A2 cells were collected and lysed as described above and protein concentration was assessed using a Pierce BCA protein assay kit (ThermoFisher Scientific). 3 aliquots of 50 µg, 20 µg, or 10 µg of CEM-A2 lysate were prepared by diluting the lysate in lysis buffer in a final volume of 30 µL, separated by SDS PAGE and analyzed by western blot as described above except the membrane was blocked with 5% BSA in PBS.

3 aliquots of 10 µg of NHE6 antibody (ThermoFisher Scientific) were diluted in 5% BSA in PBS. 20 µL of solvent was added to one aliquot to produce a solvent control. 50 µg of epitope peptide describe above was added to one aliquot of antibody (5X excess the antibody amount). 100 µg of epitope peptide was added to one aliquot of antibody (10X excess the antibody amount). The antibody aliquots were pre-incubated with the peptide or solvent for 7 hours at 4 °C with rotation. After incubation with the epitope peptide or solvent, the antibodies were added to the membranes, incubated overnight at 4 °C with rocking and stained with antibody directed against NHE6 as described above.

To ensure the epitope peptide was specific to the NHE6 antibody and would not block detection of other NHEs, the experiment was performed exactly as described above, but with the NHE8 antibody (ProteinTech).

### Quantification and statistical analysis

#### Calculation of Relative Protein Expression

For Figure 1E-I, chemiluminescent signal of the target proteins was quantified with Image Lab software (Bio-Rad) and divided by the corresponding vinculin signal to control for slight variations in total protein. The fold change of protein expression relative to GFP^-^ samples was calculated as follows for each target protein:

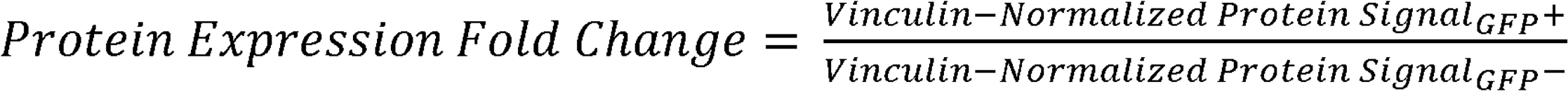

Statistical significance was determined with a One-Way ANOVA mixed effects analysis with Dunnett correction.

The same analysis and statistical test was performed for Supplemental Figure S5B and C.

#### Quantification of Changes in Transferrin^+^ Compartment Acidity

For Figure 1J, MFIs for pHrodo and AlexFluor647 were obtained with FlowJo software for the GFP^+^ and mock infected cells. The pHrodo signal was first normalized to the AlexaFluor 647 signal using the following equation to account for variations in transferrin uptake between samples:

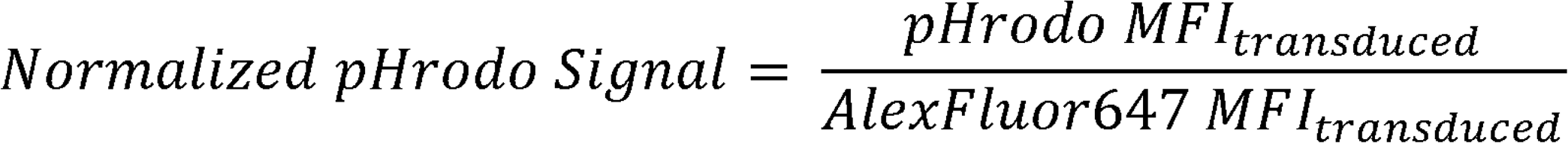

Fold change of pHrodo signal relative to mock infected cells was calculated as follows:

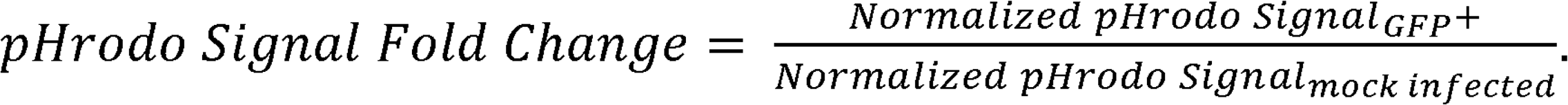

The same analysis was conducted for Figure 2H and I. Statistical significance was determined by a One-Way ANOVA mixed effects analysis with Tukey correction with GraphPad Prism software.

#### Quantification of NHE overexpression by confocal microscopy

For Figure 2B and D and Supplemental Figure S4B, cell area, integrated density, and mean grey value were obtained from Fiji software for each cell and sample of background from each image. Corrected total cell fluorescence (CTCF) was calculated as follows for each cell:

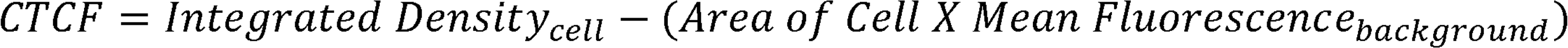

CTCF for at least 30 cells transduced with the NHE overexpressing viruses or ΔGPE NL-GI-IRES were calculated. Fluorescent signal fold change for each NHE was calculated with the following equation:

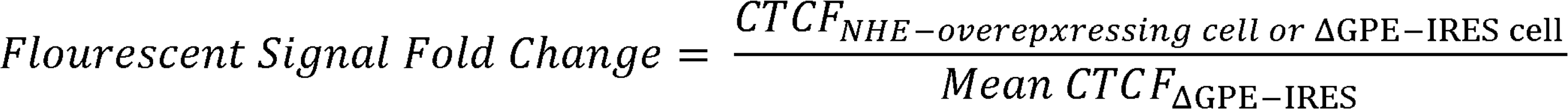

The resulting values were plotted as bar graphs and statistical significance was determined by an unpaired T-test with GraphPad Prism software.

#### Calculation of Fold Change of Nef-dependent MHC-I and CD4 downmodulation between ΔGPE-IRES-NHE6/8, ΔGPE-IRES, and ΔGPEN

As shown in Figure 5B, downmodulation of MHC-I by Nef, referred to as Nef activity, was calculated by performing flow cytometric analysis of its expression. Median fluorescent intensities (MFIs) for MHC-I expression of live GFP^-^ cells (untransduced) and GFP^+^ cells (transduced) was calculated with FlowJo data analysis software. These values were applied to the following equation:

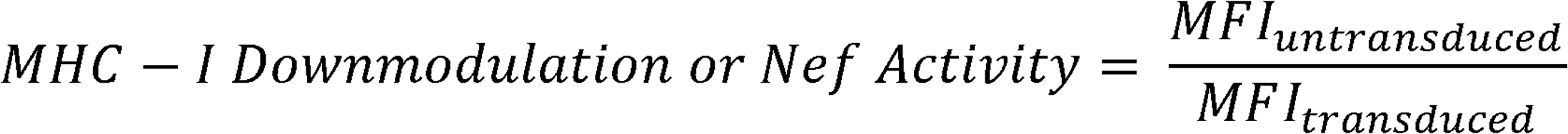

For Figures 2F, to determine if MHC-I downmodulation was impacted by NHE overexpression, its downmodulation was calculated as described above for primary CD4^+^ T cells transduced with ΔGPE-IRES-NHE6/8, ΔGPE-IRES, or ΔGPEN. The fold change in downmodulation of MHC-I for each virus compared to ΔGPE NL-GI-IRES was calculated as follows:

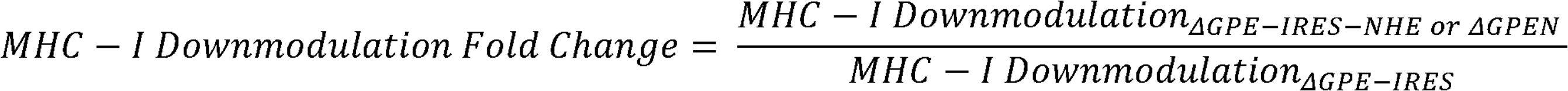

CD4 downmodulation was calculated as described for MHC-I, except that CD4 median fluorescence intensity (MFI) was utilized. For figure 2G, CD4 downmodulation fold change was calculated exactly as described for MHC-I. Statistical significance for Figure 2F and G was determined with a One-Way ANOVA mixed effects analysis with Dunnett correction. For Figure 5B, statistical significance between downmodulation of MHC-I and CD4 for ΔGPE-IRES and ΔGPE-FLAG-Nef-IRES was determined with paired T test calculated with GraphPad Prism software.

#### Object-based quantification of colocalization

To quantify colocalization between overexpressed NHE6 with Rab11, Rab5, Rab7, or TGN46, images of cells were uploaded in Imaris image analysis software and regions of interest were drawn around GFP^+^ cells with bright staining for all proteins of interest. The average diameter of puncta corresponding to all stains was measured across multiple cells for each set of images (average diameters for NHE6 were 0.7 µm, Rab11 0.7 µm, Rab5 0.7 µm, TGN46 0.6 µm). The average diameter of the puncta were used to create a spot mask for each stain and a quality threshold was set and maintained across all images from the same experimental replicate to differentiate true staining from background. Spots were considered colocalized if the distance between them was less than or equal to the sum of the radii for each stain (i.e. NHE6 and TNG46 were considered colocalized if the distance between the corresponding spots was less than or equal to 0.65 µm). For each image, the number of NHE6 spots that colocalized with the corresponding organelle marker was divided by the total number of NHE6 spots to quantify colocalization. The corresponding values were graphed as shown in Figure 3F where each point represents one image. The number of cells quantified ranged from 1-8 cells per image. Quantification of colocalization of transferrin and Rab11 or Rab5 was performed exactly as described above. Graphs are displayed in Supplemental Figure S1E. Quantification of colocalization between overexpressed NHE9 and Rab7 and NHE8 and AP-1 γ was performed exactly as described above. Graphs are displayed in Supplemental Figure S3B (NHE8 and AP-1 γ) and Supplemental Figure S4E (NHE9 and Rab7).

Quantification of colocalization between Nef and Rab11 with and without NHE6 overexpression was performed as described above except a surface mask was created over the Nef staining instead of a spots mask because Nef localizes to multiple places in the cell and the resulting staining did not produce discrete puncta. The surface detail was 0.5 µm and a quality threshold was set and maintained across all images from the same experimental replicate to distinguish between true staining and background. A spots mask was superimposed over the Rab11 staining with the diameter of the spots being set to 0.5 µm. Nef and Rab11 were considered colocalized if the average distance between the Nef surface mask and individual Rab11 spots was less than or equal to 0 µm. For each image, the total number of Rab11 spots colocalized with the Nef mask was divided by the total number of Rab11 spots and this value was graphed as shown in Figure 6G where each point represents one image. The number of cells quantified per imaged ranged from 1-9. Statistical significance was determined with an unpaired T test using GraphPad Prism software.

#### Quantification of Lysosome neutralization in response to NHE overexpression

For Figure 2J, the extent of lysosome acidification in response to NHE overexpression was evaluated by obtaining Lysotracker MFIs for GFP^+^ cells transduced with ΔGPE NL-GI-IRES-NHE6/8, ΔGPE NL-GI-IRES, or ΔGPEN from FlowJo software. Fold change of Lysotracker signal in response to NHE overexpression or ΔGPEN compared to no overexpression was calculated with the following equation:

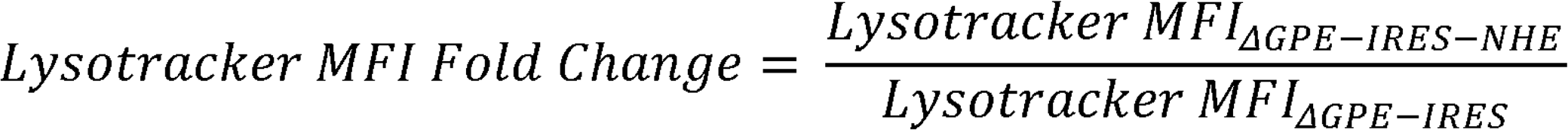

#### Calculation of CMA and Baf A1 Dose response curves and IC_50_ values for Nef Activity, Transferrin^+^ Compartment Neutralization, and Lysotracker

At every concentration of CMA or Baf A1 and solvent tested, MHC-I downmodulation was calculated as described above. MHC-I downmodulation in the presence of CMA or Baf A1 was compared to solvent using the following equation for each concentration of CMA or Baf A1 tested:

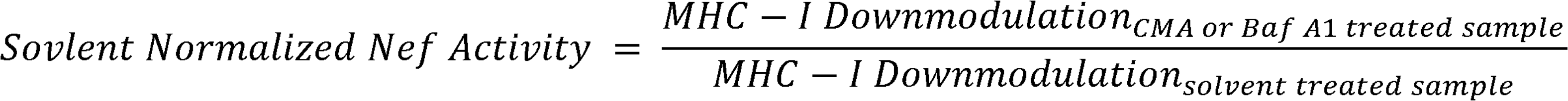

The resulting data were plotted to produce dose-response curves seen in Figure 4A, B, D, E, and G. The amount of CMA or Baf A1 required to inhibit 50% of Nef activity (CMA IC_50_ or Baf A1 IC_50,_ Nef Activity), as seen in Figure 4C, F, and H, were calculated with GraphPad Prism software.

The impact of CMA or Baf A1 on lysosome acidification was calculated exactly like that for Nef Activity, except using Lysotracker MFIs for GFP^-^ cells instead of MHC-I downmodulation in the numerator and denominator (Figure 4A and D).

Transferrin^+^ compartmental neutralization by CMA or Baf A1 was assessed by flow cytometric analysis. The pHrodo signal was normalized to the AF-647 signal using the normalized pHrodo signal equation above. At every concentration of CMA, Baf A1, or solvent tested, the normalized pHrodo signal of the GFP^+^ CMA or Baf A1 treated cells was compared to the solvent treated cells using the following equation:

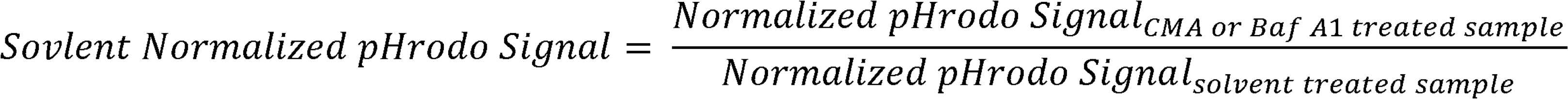

The resulting data points were plotted to produce the dose-response curve shown in Figure 4B, E and G. The amount of CMA or Baf A1 required to reduce the pHrodo signal by 50% (CMA or Baf A1 IC_50_, pHrodo-Transferrin MFI) was obtained from GraphPad Prism software. Further, the IC_50_s for Nef activity, pHrodo-transferrin MFI, and Lysotracker MFI were compared as shown in Figure 4C and 4F. For Figure 4C and 4F, statistical significance was determined with a One-Way ANOVA mixed effects analysis with Dunnet correction calculated with GraphPad Prism software. A paired T test was used to determine statistical significance Figure 4H.

#### Quantification of protein complex formation by Nef during NHE6 overexpression

Western blots of co-immunoprecipitation experiments of CEM-A2 lysates transduced with ΔGPE-FLAG-Nef-IRES, ΔGPE-FLAG-Nef-IRES-NHE6, or mock transduced were imaged on an iBright Imaging System (ThermoFisher Scientific). Intensity of protein bands was obtained with ImageStudio software (LI-COR). To account for slight variations in the amount of Nef pulled down between samples, chemiluminescent signals of the target proteins were first normalized to Nef intensities using the following equation:

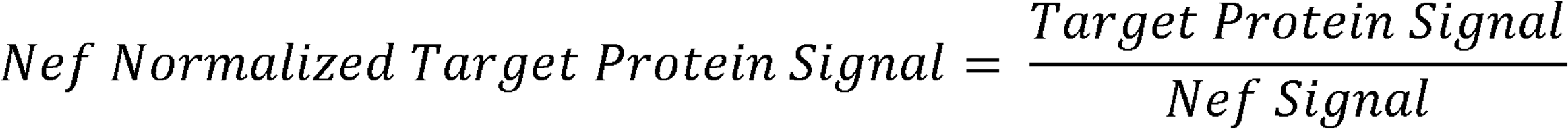

The fold change of protein bound to Nef between ΔGPE-FLAG-Nef-IRES and ΔGPE-FLAG-Nef-IRES-NHE6 was then calculated as follows:

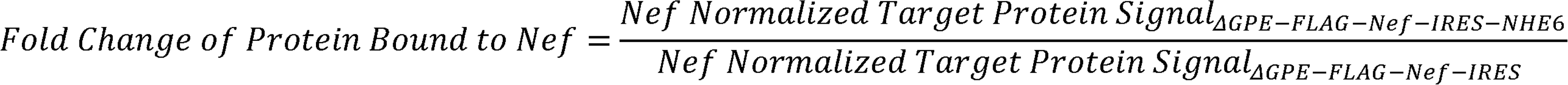

This was graphed as shown in Figure 5E. Statistically significant changes binding of each target protein to Nef were evaluated with a paired T test using GraphPad Prism software.

#### Quantification of Nef Present in Rab11^+^ Compartments with and without NHE6 Overexpression

Western blots of the contents of precipitated intact Rab11^+^ compartments were imaged on a Chemidoc Imaging system (Bio-Rad). Intensity of protein bands was obtained with Image Lab software (Bio-Rad). To account for slight variations in the amount of Rab11 pulled down between samples, Nef signal was first normalized to the corresponding Rab11 signal with the following equation:

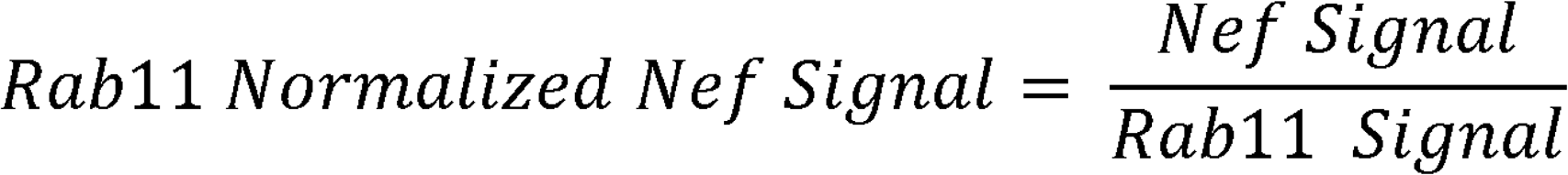

The fold change of the amount of Nef found in isolated Rab11^+^ compartments relative to transduction with ΔGPE-NL-GI-IRES was calculated as follows:

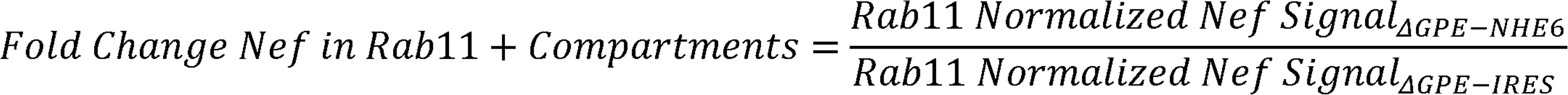

The resulting values were graphed as shown in Figure 6E. Statistical significance was determined with a Paired T Test calculated with GraphPad Prism software.

## REFERENCES

1. Caffrey, M., and Lavie, A. (2021). pH-Dependent Mechanisms of Influenza Infection Mediated by Hemagglutinin. Front Mol Biosci 8, 777095. 10.3389/fmolb.2021.777095.

2. Doyle, T., Moncorgé, O., Bonaventure, B., Pollpeter, D., Lussignol, M., Tauziet, M., Apolonia, L., Catanese, M.-T., Goujon, C., and Malim, M.H. (2018). The interferon-inducible isoform of NCOA7 inhibits endosome-mediated viral entry. Nat Microbiol 3, 1369–1376. 10.1038/s41564-018-0273-9.

3. Khan, H., Winstone, H., Jimenez-Guardeño, J.M., Graham, C., Doores, K.J., Goujon, C., Matthews, D.A., Davidson, A.D., Rihn, S.J., Palmarini, M., et al. (2021). TMPRSS2 promotes SARS-CoV-2 evasion from NCOA7-mediated restriction. PLOS Pathogens 17, e1009820. 10.1371/journal.ppat.1009820.

4. Miyauchi, K., Kim, Y., Latinovic, O., Morozov, V., and Melikyan, G.B. (2009). HIV Enters Cells via Endocytosis and Dynamin-Dependent Fusion with Endosomes. Cell 137, 433–444. 10.1016/j.cell.2009.02.046.

5. Nakamura, N., Tanaka, S., Teko, Y., Mitsui, K., and Kanazawa, H. (2005). Four Na+/H+ Exchanger Isoforms Are Distributed to Golgi and Post-Golgi Compartments and Are Involved in Organelle pH Regulation*. Journal of Biological Chemistry 280, 1561–1572. 10.1074/jbc.M410041200.

6. Ohgaki, R., Matsushita, M., Kanazawa, H., Ogihara, S., Hoekstra, D., and van IJzendoorn, S.C.D. (2010). The Na+/H+ Exchanger NHE6 in the Endosomal Recycling System Is Involved in the Development of Apical Bile Canalicular Surface Domains in HepG2 Cells. Mol Biol Cell 21, 1293–1304. 10.1091/mbc.E09-09-0767.

7. Xinhan, L., Matsushita, M., Numaza, M., Taguchi, A., Mitsui, K., and Kanazawa, H. (2011). Na+/H+ exchanger isoform 6 (NHE6/SLC9A6) is involved in clathrin-dependent endocytosis of transferrin. Am J Physiol Cell Physiol 301, C1431–1444. 10.1152/ajpcell.00154.2011.

8. Painter, M.M., Zimmerman, G.E., Merlino, M.S., Robertson, A.W., Terry, V.H., Ren, X., McLeod, M.R., Gomez-Rodriguez, L., Garcia, K.A., Leonard, J.A., et al. (2020). Concanamycin A counteracts HIV-1 Nef to enhance immune clearance of infected primary cells by cytotoxic T lymphocytes. Proceedings of the National Academy of Sciences 117, 23835–23846. 10.1073/pnas.2008615117.

9. Roeth, J.F., Williams, M., Kasper, M.R., Filzen, T.M., and Collins, K.L. (2004). HIV-1 Nef disrupts MHC-I trafficking by recruiting AP-1 to the MHC-I cytoplasmic tail. J Cell Biol 167, 903–913. 10.1083/jcb.200407031.

10. Kasper, M.R., and Collins, K.L. (2003). Nef-Mediated Disruption of HLA-A2 Transport to the Cell Surface in T Cells. Journal of Virology 77, 3041–3049. 10.1128/jvi.77.5.3041-3049.2003.

11. Schaefer, M.R., Wonderlich, E.R., Roeth, J.F., Leonard, J.A., and Collins, K.L. (2008). HIV-1 Nef Targets MHC-I and CD4 for Degradation Via a Final Common β-COP–Dependent Pathway in T Cells. PLOS Pathogens 4, e1000131. 10.1371/journal.ppat.1000131.

12. Doyle, T., Moncorgé, O., Bonaventure, B., Pollpeter, D., Lussignol, M., Tauziet, M., Apolonia, L., Catanese, M.-T., Goujon, C., and Malim, M.H. (2019). Author Correction: The interferon-inducible isoform of NCOA7 inhibits endosome-mediated viral entry. Nat Microbiol 4, 539. 10.1038/s41564-019-0366-0.

13. Brett, C.L., Wei, Y., Donowitz, M., and Rao, R. (2002). Human Na+/H+ exchanger isoform 6 is found in recycling endosomes of cells, not in mitochondria. American Journal of Physiology-Cell Physiology 282, C1031–C1041. 10.1152/ajpcell.00420.2001.

14. Ilie, A., Boucher, A., Park, J., Berghuis, A.M., McKinney, R.A., and Orlowski, J. (2020). Assorted dysfunctions of endosomal alkali cation/proton exchanger SLC9A6 variants linked to Christianson syndrome. J Biol Chem 295, 7075–7095. 10.1074/jbc.RA120.012614.

15. Ilie, A., Gao, A.Y.L., Reid, J., Boucher, A., McEwan, C., Barrière, H., Lukacs, G.L., McKinney, R.A., and Orlowski, J. (2016). A Christianson syndrome-linked deletion mutation (Δ287ES288) in SLC9A6 disrupts recycling endosomal function and elicits neurodegeneration and cell death. Mol Neurodegener 11, 63. 10.1186/s13024-016-0129-9.

16. Ilie, A., Weinstein, E., Boucher, A., McKinney, R.A., and Orlowski, J. (2014). Impaired posttranslational processing and trafficking of an endosomal Na+/H+ exchanger NHE6 mutant (Δ370WST372) associated with X-linked intellectual disability and autism. Neurochemistry International 73, 192–203. 10.1016/j.neuint.2013.09.020.

17. Pruett, B.S., Pinner, A.L., Kim, P., and Meador-Woodruff, J.H. (2023). Altered distribution and localization of organellar Na+/H+ exchangers in postmortem schizophrenia dorsolateral prefrontal cortex. Transl Psychiatry 13, 1–10. 10.1038/s41398-023-02336-2.

18. Mayle, K.M., Le, A.M., and Kamei, D.T. (2012). The Intracellular Trafficking Pathway of Transferrin. Biochim Biophys Acta 1820, 264–281. 10.1016/j.bbagen.2011.09.009.

19. Wonderlich, E.R., Leonard, J.A., Kulpa, D.A., Leopold, K.E., Norman, J.M., and Collins, K.L. (2011). ADP Ribosylation Factor 1 Activity Is Required To Recruit AP-1 to the Major Histocompatibility Complex Class I (MHC-I) Cytoplasmic Tail and Disrupt MHC-I Trafficking in HIV-1-Infected Primary T Cells. Journal of Virology 85, 12216–12226. 10.1128/jvi.00056-11.

20. Park, H., Hundley, F.V., Yu, Q., Overmyer, K.A., Brademan, D.R., Serrano, L., Paulo, J.A., Paoli, J.C., Swarup, S., Coon, J.J., et al. (2022). Spatial snapshots of amyloid precursor protein intramembrane processing via early endosome proteomics. Nat Commun 13, 6112. 10.1038/s41467-022-33881-x.

21. Cotter, K., Stransky, L., McGuire, C., and Forgac, M. (2015). Recent Insights into the Structure, Regulation, and Function of the V-ATPases. Trends Biochem Sci 40, 611–622. 10.1016/j.tibs.2015.08.005.

22. Trombetta, E.S., Ebersold, M., Garrett, W., Pypaert, M., and Mellman, I. (2003). Activation of Lysosomal Function During Dendritic Cell Maturation. Science 299, 1400–1403. 10.1126/science.1080106.

23. Kasper, M.R., and Collins, K.L. (2003). Nef-Mediated Disruption of HLA-A2 Transport to the Cell Surface in T Cells. Journal of Virology 77, 3041–3049. 10.1128/jvi.77.5.3041-3049.2003.

24. Schaefer, M.R., Wonderlich, E.R., Roeth, J.F., Leonard, J.A., and Collins, K.L. (2008). HIV-1 Nef Targets MHC-I and CD4 for Degradation Via a Final Common β-COP–Dependent Pathway in T Cells. PLOS Pathogens 4, e1000131. 10.1371/journal.ppat.1000131.

25. Wonderlich, E.R., Leonard, J.A., Kulpa, D.A., Leopold, K.E., Norman, J.M., and Collins, K.L. (2011). ADP Ribosylation Factor 1 Activity Is Required To Recruit AP-1 to the Major Histocompatibility Complex Class I (MHC-I) Cytoplasmic Tail and Disrupt MHC-I Trafficking in HIV-1-Infected Primary T Cells. Journal of Virology 85, 12216–12226. 10.1128/jvi.00056-11.

26. Roeth, J.F., Williams, M., Kasper, M.R., Filzen, T.M., and Collins, K.L. (2004). HIV-1 Nef disrupts MHC-I trafficking by recruiting AP-1 to the MHC-I cytoplasmic tail. J Cell Biol 167, 903–913. 10.1083/jcb.200407031.

27. Kuo, R.-L., Lin, Y.-H., Wang, R.Y.-L., Hsu, C.-W., Chiu, Y.-T., Huang, H.-I., Kao, L.-T., Yu, J.-S., Shih, S.-R., and Wu, C.-C. (2015). Proteomics Analysis of EV71-Infected Cells Reveals the Involvement of Host Protein NEDD4L in EV71 Replication. J. Proteome Res. 14, 1818–1830. 10.1021/pr501199h.

28. Fraisier, C., Koraka, P., Belghazi, M., Bakli, M., Granjeaud, S., Pophillat, M., Lim, S.M., Osterhaus, A., Martina, B., Camoin, L., et al. (2014). Kinetic Analysis of Mouse Brain Proteome Alterations Following Chikungunya Virus Infection before and after Appearance of Clinical Symptoms. PLOS ONE 9, e91397. 10.1371/journal.pone.0091397.

29. Baik, S.Y., Yun, H.S., Lee, H.J., Lee, M.H., Jung, S.E., Kim, J.W., Jeon, J.P., Shin, Y.K., Rhee, H.S., Kimm, K.C., et al. (2007). Identification of stathmin 1 expression induced by Epstein–Barr virus in human B lymphocytes. Cell Prolif 40, 268–281. 10.1111/j.1365-2184.2007.00429.x.

30. Muramoto, Y., Shoemaker, J.E., Le, M.Q., Itoh, Y., Tamura, D., Sakai-Tagawa, Y., Imai, H., Uraki, R., Takano, R., Kawakami, E., et al. (2014). Disease Severity Is Associated with Differential Gene Expression at the Early and Late Phases of Infection in Nonhuman Primates Infected with Different H5N1 Highly Pathogenic Avian Influenza Viruses. J Virol 88, 8981–8997. 10.1128/JVI.00907-14.

31. Prasad, H. (2021). Protons to Patients: targeting endosomal Na+ /H+ exchangers against COVID-19 and other viral diseases. FEBS J 288, 5071–5088. 10.1111/febs.16163.

32. Slonchak, A., Clarke, B., Mackenzie, J., Amarilla, A.A., Setoh, Y.X., and Khromykh, A.A. (2019). West Nile virus infection and interferon alpha treatment alter the spectrum and the levels of coding and noncoding host RNAs secreted in extracellular vesicles. BMC Genomics 20, 474. 10.1186/s12864-019-5835-6.

33. Lu, X., Yu, H., Liu, S.H., Brodsky, F.M., and Peterlin, B.M. (1998). Interactions between HIV1 Nef and vacuolar ATPase facilitate the internalization of CD4. Immunity 8, 647–656. 10.1016/s1074-7613(00)80569-5.

