## Supplementary material for "Virus-induced vesicular acidification enhances HIV immune evasion": Virus-induced vesicular acidification enhances HIV immune evasion._SI

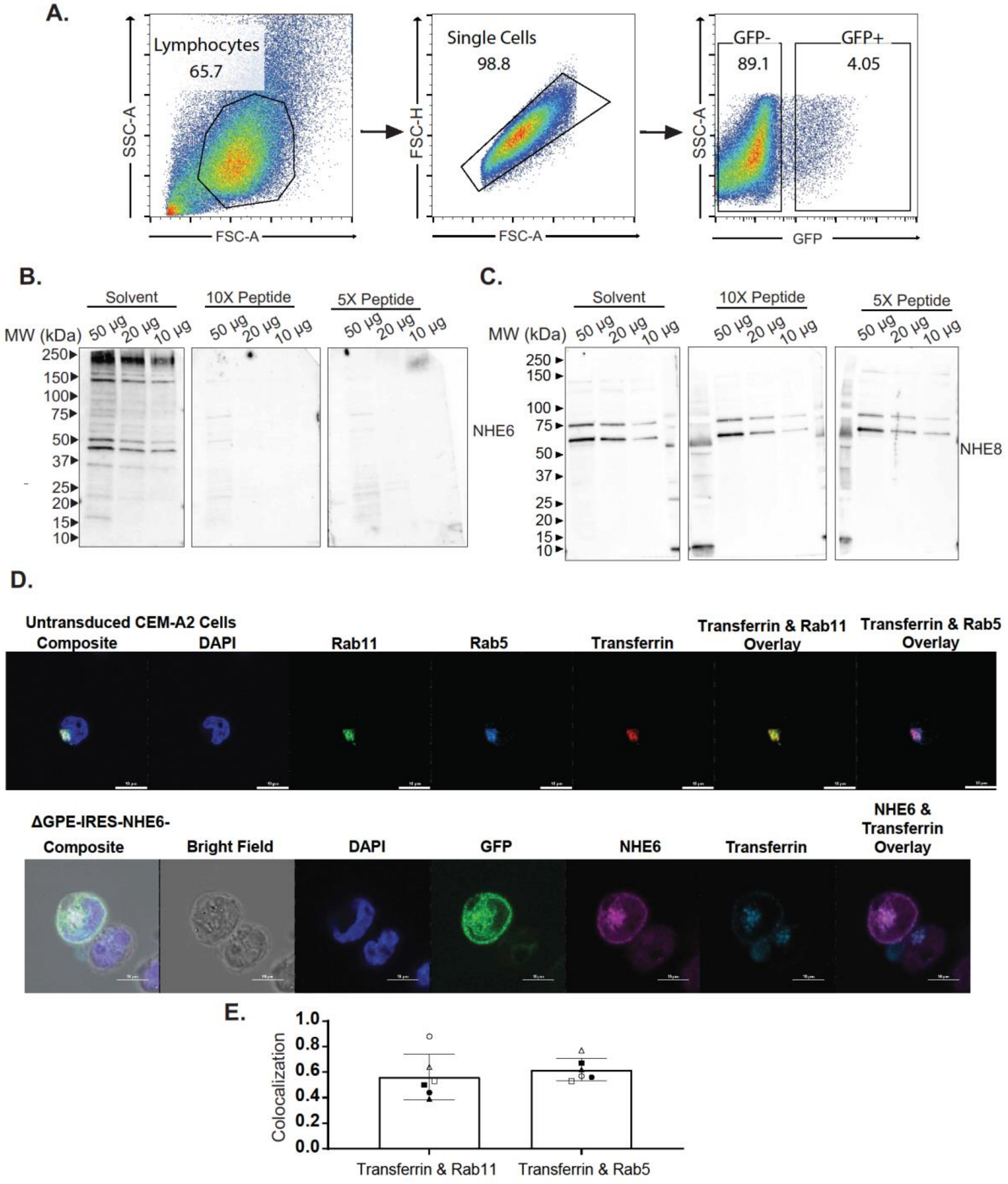


**Supplemental Figure S1. Characterization of NHE6 expression.** Related to Figure 1. (A) General gating strategy for flow cytometry experiments. (B) NHE6 peptide epitope competitively blocks NHE6 antibody binding. Western blot analysis of the indicated amounts of CEM-A2 cell lysates incubated with primary NHE6 antibody that was preincubated with solvent or a peptide epitope of the NHE6 antibody at 10X excess of the antibody amount (100 μg of peptide) or 5X excess the antibody amount (50 μg of peptide). (C) NHE6 peptide epitope does not inhibit binding of non-NHE6 antibodies. Western blot analysis of the indicated amounts of CEM-A2 cell lysates incubated with primary NHE8 antibody that was preincubated with solvent or a peptide epitope of the NHE6 antibody as described in (B). (D) NHE6 co-localizes with transferrin. Representative confocal fluorescence microscopy images of untransduced CEM-A2 cells incubated with transferrin AF-647 for 15 minutes (time point of our assay) then stained for Rab5 and Rab11 (top panel). Confocal fluorescence microscopy images of CEM-A2 cells transduced with ΔGPE-IRES-NHE6 and incubated with transferrin AF-647 for 15 minutes, then stained for NHE6 (bottom). (E) Transferrin colocalizes with Rab5 and Rab11. Summary graph of colocalization of transferrin with Rab5 and Rab11. Colocalization was quantified as described in Figure 3F. A spots mask was assigned to transferrin, Rab5, and Rab11. The number of transferrin spots colocalized with Rab5 or Rab11 was divided by the total number of transferrin spots and graphed. Each point represents one image. Each symbol represents quantification of colocalization of transferrin with Rab5 or Rab11 from the same image. A total of 6 images were analyzed. The number of cells analyzed per imaged ranged from 2-4. 1 biological replicate was performed.

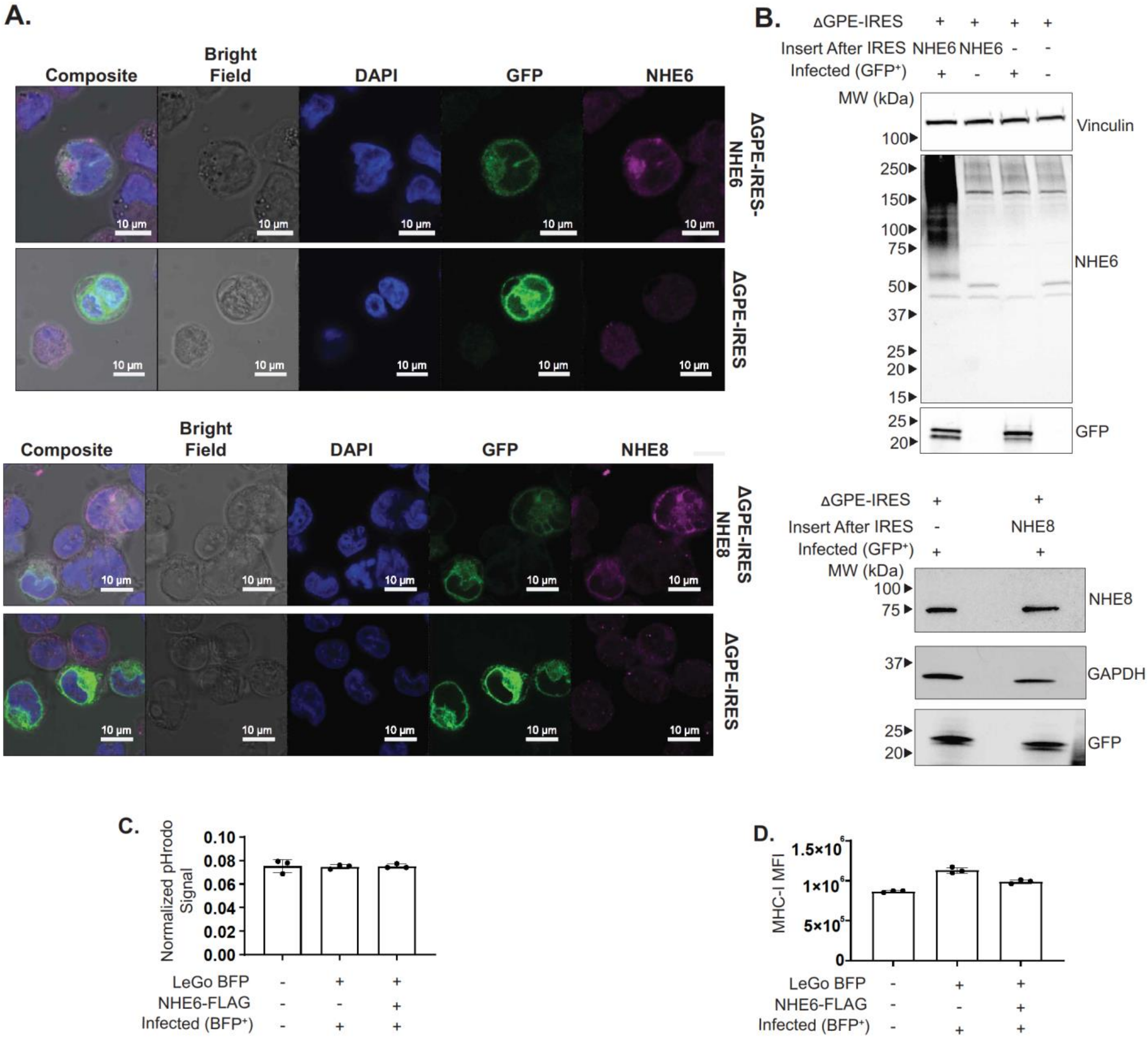


**Supplemental Figure S2**. Characterization of NHE6 and 8 expression. Related to Figure 2. (A) Full confocal microscopy images from Figure 2A, and C showing DAPI staining, bright field, and composite images. (B) Confirmation of NHE6 and NHE8 overexpression by western blot. Western blot analysis of lysates from CEM-A2 cells 48 hr post transduction with the indicated viruses and subjected to FACS analysis to isolate GFP+ (transduced) or GFP- (untransduced) cells and probed for the indicated NHEs. (C) NHE6 overexpression does not alter the pH of transferrin+ compartments in uninfected cells. Summary graph of transferrin+ compartment acidity of primary CD4+ T cells 3 days post transduction with the indicated viruses that either overexpress FLAG-tagged NHE6 (LeGo-BFP-NHE6-FLAG) or empty vector (LeGo-BFP). Cells were stained with pHrodo-transferrin and AF-647 transferrin. pHrodo transferrin MFIs for BFP+ (transduced) and BFP- (untransduced) cells were normalized to the corresponding AF-647 transferrin MFIs and graphed. One biological replicate was performed in technical triplicate. (D) NHE6 overexpression does not alter surface MHC-I expression in uninfected cells. Summary graph of surface MHC-I expression of primary CD4+ T cells 3 days post transduction with the indicated viruses. MHC-I MFIs for BFP+ (transduced) and BFP- (untransduced) cells were obtained and graphed. One biological replicate was performed in technical triplicate

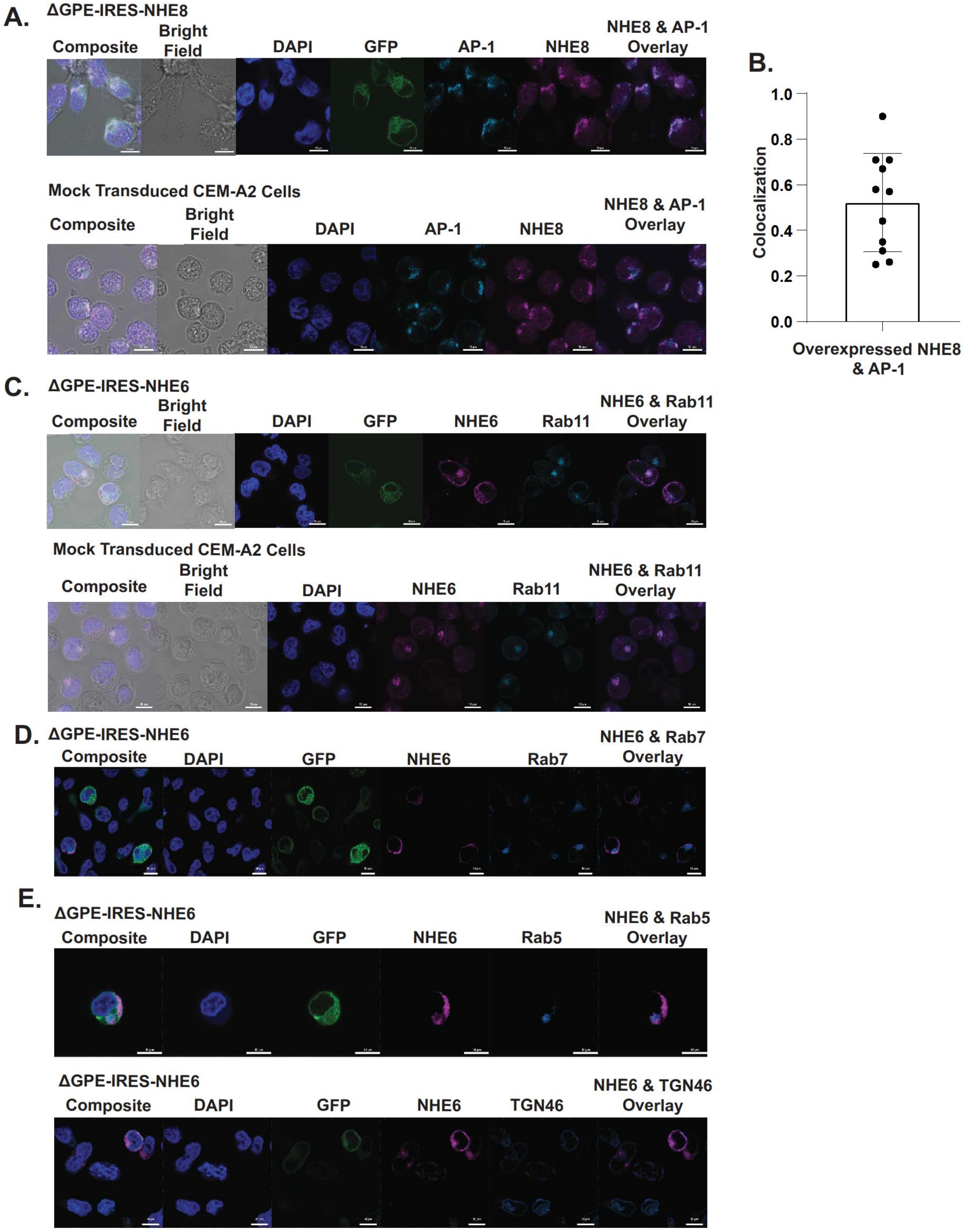

A.
ΔGPE-IRES-NHE8
Composite
Bright Field
DAPI
GFP
AP-1
NHE8
NHE8 & AP-1 Overlay
Mock Transduced CEM-A2 Cells
Composite
Bright Field
DAPI
AP-1
NHE8
NHE8 & AP-1 Overlay
B.
1.0
0.8
0.6
0.4
0.2
0.0
Colocalization
Overexpressed NHE8 & AP-1
C.
ΔGPE-IRES-NHE6
Composite
Bright Field
DAPI
GFP
NHE6
Rab11
NHE6 & Rab11 Overlay
Mock Transduced CEM-A2 Cells
Composite
Bright Field
DAPI
NHE6
Rab11
NHE6 & Rab11 Overlay
D.
ΔGPE-IRES-NHE6
Composite
DAPI
GFP
NHE6
Rab7
NHE6 & Rab7 Overlay
E.
ΔGPE-IRES-NHE6
Composite
DAPI
GFP
NHE6
Rab5
NHE6 & Rab5 Overlay
ΔGPE-IRES-NHE6
Composite
DAPI
GFP
NHE6
TGN46
NHE6 & TGN46 Overlay

**Supplemental Figure S3.** Characterization of colocalization of NHE6 and NHE8 in CEM-A2 cells. Related to Figure 3. (A) HIV infection does not alter AP-1 staining. (Top panel) Full confocal images from Figure 3A including composite, bright field, and DAPI staining. (Bottom panel) NHE8 and AP-1 staining of mock transduced CEM-A2 cells. (B) Summary graph of quantification of colocalization between overexpressed NHE8 and AP-1 γ. Colocalization was quantified as described in Figure 3F. A spots mask was assigned for NHE8 and AP-1 γ staining. The number of NHE8 spots that colocalized with AP-1 γ was divided by the total number of NHE8 spots and graphed. Each point represents 1 image. A total of 12 images were analyzed. The number of cells quantified per image ranged from 1-4. 1 biological replicate was performed. (C) (Top panel) Full confocal images from Figure 3B including composite, bright field, and DAPI staining. (Bottom panel) staining of NHE6 and Rab11 in mock transduced CEM-A2 cells. (D) Full confocal images from Figure 3C including composite and DAPI staining. (E) Full confocal images from Figure 3D including composite and DAPI staining. (F) Full confocal images from Figure 3E including composite and DAPI staining.

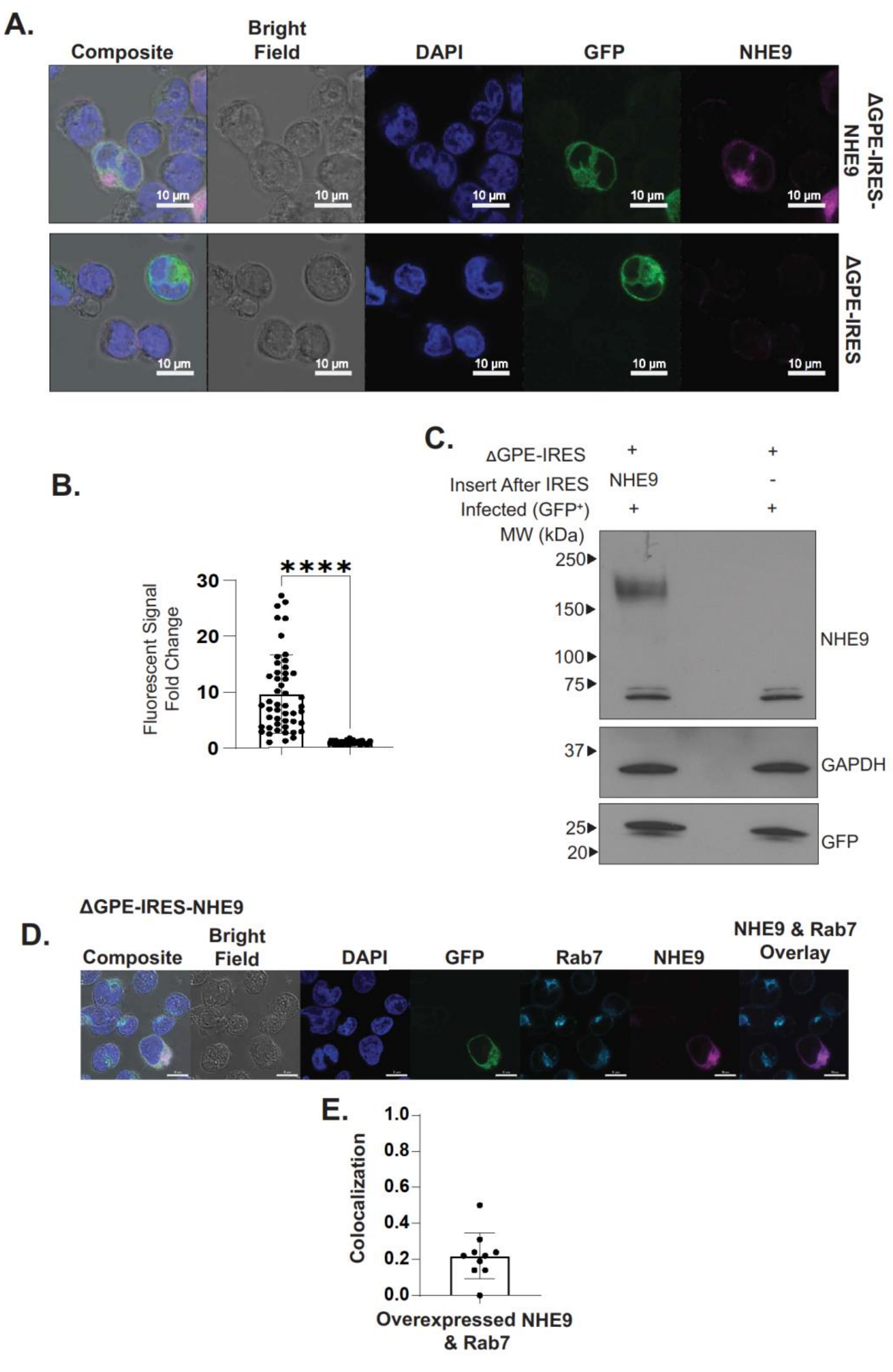


**Supplemental Figure S4.** Overexpressed NHE9 does not colocalize with Rab7 in CEM-A2 cells. Related to Figure 3. (A) Representative confocal fluorescence microscopy images of CEM-A2 cells 48 hr post transduction with the indicated viruses and stained for NHE9 and Rab7. (B) Summary graph of quantification of expression of NHE8 48 hr post transduction of CEM-A2 cells with the indicated viruses. CTCF and fold change were calculated as described in Figure 2B. At least 30 cells were imaged. (C) Western blot analysis of CEM-A2 lysates 48 hr post transduction with the indicated viruses and sorted for infected (GFP+) cells. (D) Representative confocal fluorescence microscopy images of CEM-A2 cells 48 hr post transduction with the indicated virus then stained for NHE9 and Rab7. (E) Summary graph of quantification of colocalization between overexpressed NHE9 and Rab7. Colocalization was quantified as described in Figure 3F. A spots mask was assigned to NHE9 and Rab7 staining. The number of NHE9 spots colocalized with Rab7 was divided by the total number of NHE9 spots and graphed. Each point represents 1 image. 10 images were analyzed. The number of cells analyzed per image ranged from 1-3. 1 biological replicate was performed.

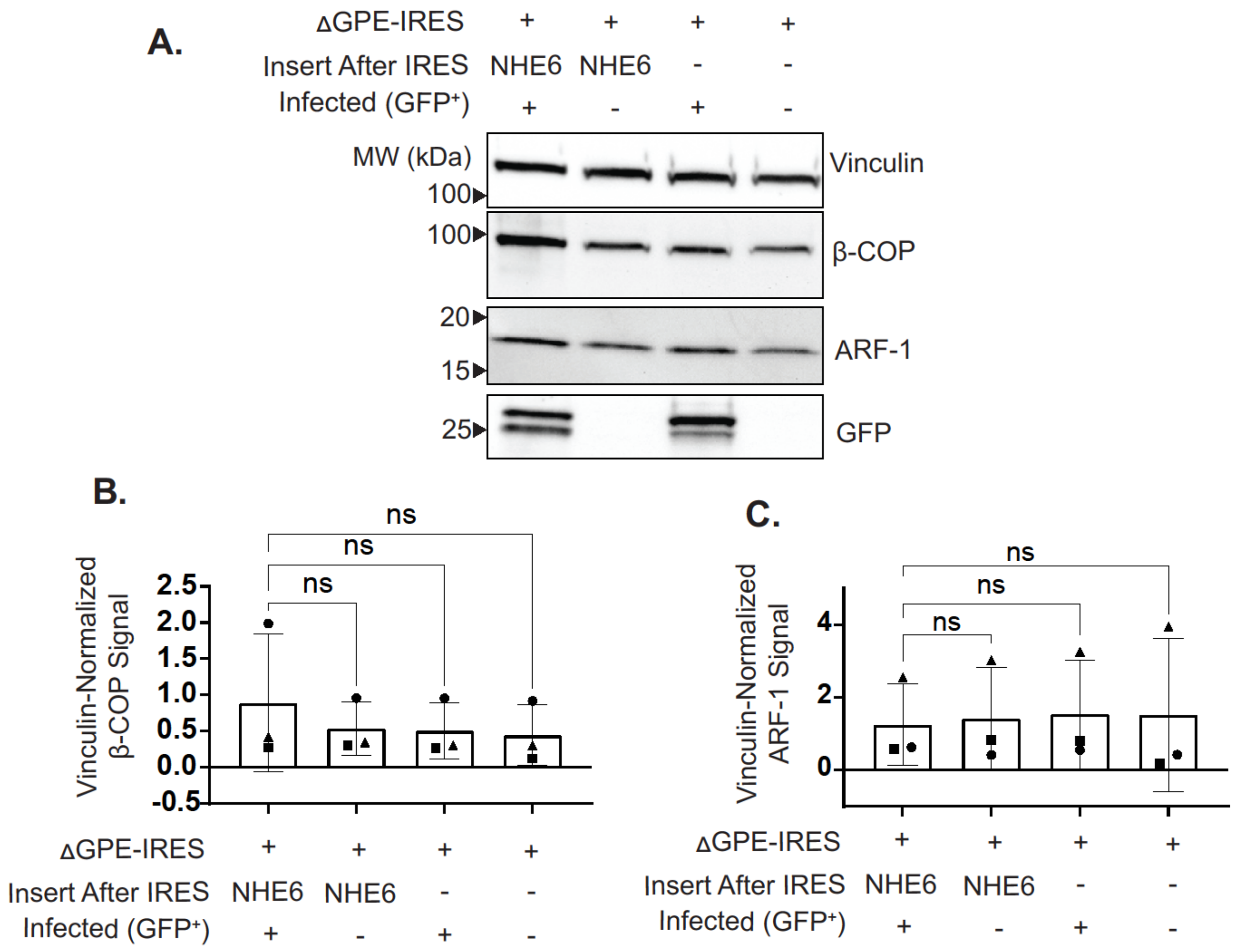


**Supplemental Figure S5.** NHE6 overexpression does not alter expression of β-COP or ARF-1. Related to Figure 5. (A) Western blot analysis of lysates from CEM-A2 cells 48 hr post transduction with the indicated viruses and subjected to FACS analysis to isolate the GFP+ (transduced) or GFP- (untransduced) populations. (B) Summary graph of β-COP protein expression from western blot analysis described in (A). β-COP signal was normalized to the corresponding vinculin signal and graphed. 3 biological replicates were performed. (C) Summary graph of ARF-1 protein expression from western blot analysis described in (A). ARF-1 signal was normalized to the corresponding vinculin signal and graphed. 3 biological replicates were performed. Statistical significance for (D) and (E) was determined with a One Way ANOVA mixed-effects analysis with Dunnett correction. * $p < 0.05$, ** $p < 0.01$, *** $p < 0.001$, **** $p < 0.0001$

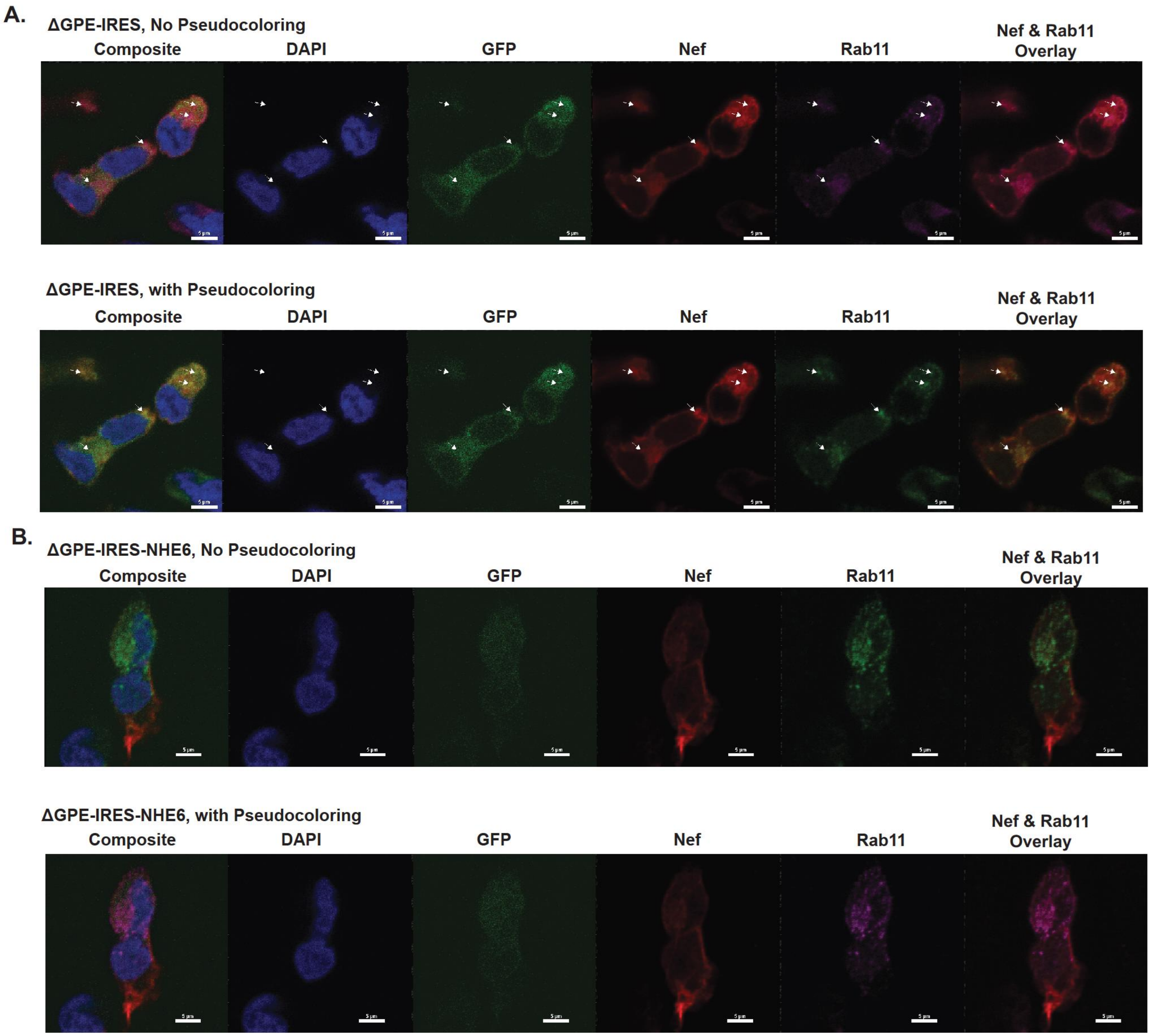


**Supplemental Figure S6.** Characterization of colocalization between Nef and Rab11 with and without NHE6 overexpression. Related to Figure 6. (A) Full confocal images without (top panel) or with (bottom panel) pseudocoloring of Rab11 from Figure 6F (top panel) including composite and DAPI staining. (B) Full confocal images without (top panel) or with (bottom panel) pseudocoloring of Rab11 from Figure 6F (bottom panel) including composite and DAPI staining.
